# Fast and accurate modeling of TCR-peptide-MHC complexes using tFold-TCR

**DOI:** 10.1101/2025.01.12.632367

**Authors:** Fandi Wu, Yu Zhao, Yang Xiao, Chenchen Qin, Fang Wang, Zihan Wu, Long-Kai Huang, Xiao Liu, Jiangning Song, Bing He, Jamie Rossjohn, Jianhua Yao

## Abstract

Alpha-beta T cell receptor (***αβ***TCR) recognition of peptide-major histocompatibility complexes (pMHCs) is a cornerstone of the adaptive immune system. Fast and accurate modeling of TCR-pMHC structures is crucial for understanding TCR recognition of pMHCs at the molecular level, which is essential for the development of TCR-based therapeutics and vaccines. Despite significant interest, this challenge remains unresolved due to the diversity of TCR-pMHC interactions and limited structural data. Here, we present tFold-TCR, a high-throughput, end-to-end universal model for predicting three-dimensional (3D) atomic-level structures of TCR-pMHC complexes, capable of predicting TCRs of different classes and MHC structures from diverse systems. tFold-TCR leverages a specially trained, protein-protein interaction-sensitive large protein language model to extract intra- and inter-chain residue contact information and evolutionary relationships, bypassing the need for multiple sequence alignment (MSA) searches. It also features innovative structure prediction and flexible docking modules to enhance accuracy, particularly for interacting contacts. Compared to existing methods, including AlphaFold-3, tFold-TCR demonstrates a 30.7% increase in prediction success rate evaluated by DockQ and is over 25 times faster. These advancements enable large-scale structural characterization of TCRs and their interactions with pMHCs. Utilizing this capability, we constructed TCRStructDB, the largest database of TCR-pMHC structures to date, encompassing 2.2 million TCRs, 0.8 million pMHCs, and 45,000 TCR-pMHC complexes. TCRStructDB provides unprecedented insights into one of the most diverse receptor-ligand interactions in biology.

## 1 Introduction

The immune system must effectively respond to a wide array of threats to provide host protection [1–3]. The adaptive immune system, in particular, is responsible for the specific molecular recognition of a vast spectrum of foreign antigens. In cell-mediated adaptive immunity against pathogens and cancers, this recognition is mediated by the interaction between alpha-beta T cell antigen receptors (*αβ*TCRs) and antigenic peptides presented by major histocompatibility complex class I (MHC-I) or class II (MHC-II) molecules [1, 4]. TCRs consist of two distinct chains: alpha and beta chains for *αβ*TCRs, and gamma and delta chains for *γδ*TCRs. The specificity and diversity of TCRs primarily originate from the six complementarity-determining regions (CDRs), three in each chain, which simultaneously engage both the peptide and the MHC molecule. Somatic gene rearrangement, along with nucleotide insertions and deletions within the CDR3 regions, generates a TCR repertoire in each individual exhibiting remarkable clonal and structural diversity, encompassing over 10^11^ distinct receptors before thymic selection [5]. This diversity equips the human body to target a vast array of distinct antigens with high specificity, making TCRs promising candidates for novel therapeutics [6–8]. TCR-based cellular [6, 9] and soluble [10] therapeutics have shown promising results in preclinical studies and early-phase clinical trials, particularly for hematological malignancies and solid tumors refractory to conventional therapies [11].

The three-dimensional (3D) structure ultimately determines the function and activity of a protein [12]. Understanding the structural basis of TCR-pMHC complexes is essential for elucidating TCR specificity, rationally designing TCR affinity, and its implications in vaccine design, autoimmunity, and cancer therapies [1, 3, 11]. Current experimental methods for structure determination, such as cryoelectron microscopy (cryoEM), X-ray crystallography and nuclear magnetic resonance spectroscopy (NMR), are expensive and time-consuming. The extensive diversity and cross-reactivity of TCRs pose challenges for large-scale experimental studies. Additionally, the relatively low affinity of TCR-pMHC interactions, especially for autoimmune TCRs, complicates structural determination. The flexibility of CDR3 loops often results in regions that are not accurately modeled or are absent in the structures. While the number of paired TCR sequences being determined is growing exponentially, the TCR-pMHC structural database is increasing only incrementally, highlighting an unmet need to close this gap. Consequently, rapid and accurate computational modeling methods are crucial for investigating TCR-pMHC interactions at the molecular level.

Conventional algorithms for modeling unliganded TCRs [4, 13, 14] and TCR-pMHC complexes [15, 16] primarily utilize template-based modeling combined with energy minimization. These approaches often face challenges due to the scarcity of templates, the flexibility and diversity of TCR CDRs, and the wide range of TCR-pMHC docking orientations. Recently, deep learning-based structure prediction methods, particularly AlphaFold [17], have demonstrated remarkable success in predicting structures of monomeric and multimeric proteins [18, 19]. Methods such as TCRdock [20] and TCRmodel2 [21] have been developed based on AlphaFold-Multimer [18] to improve TCR-pMHC complex structure prediction. These methods have achieved high accuracy in predicting protein complexes. However, these methods rely on large-scale multiple-sequence alignments (MSAs) and are slow, limiting their potential as high-throughput tools.

Protein language models (PLMs) [22–26] have been widely applied to tasks such as protein function prediction and sequence design. These models learn intrinsic interactions within protein sequences through Masked Language Models. PLM-based structure prediction methods [25, 27] bypass the time-consuming MSA search process but are primarily used for monomer protein structure prediction. Some efforts [28] have been made to adapt PLM-based methods for multimer prediction. However, except for specific sub-problems like antibodies [29, 30], their prediction accuracy falls significantly short compared to MSA-based models.

High-precision structure prediction software, such as AlphaFold3 and ESMFold, has enabled the construction of large-scale structural databases. AlphaFold DB [31] and the ESM Metagenomic Atlas [25] have predicted and organized the structures of the vast majority of sequences in UniProt [32] and MGnify [33]. Recent studies [34, 35] have filled gaps in structural databases concerning viral proteins. However, existing structural databases primarily focus on monomeric proteins because current structure prediction software struggles with protein-protein interactions, such as antibody-antigen and TCR-pMHC complexes. This lack of complex structure databases poses significant challenges for structural biologists, as understanding protein-protein interactions is crucial for elucidating many biological processes and mechanisms. Additionally, predicting complex structures requires significant time to search for MSAs across multiple chains.

In this paper, we introduce tFold-TCR, a PLM-based end-to-end method for 3D atomic-level structure prediction of TCR-pMHC complexes. tFold-TCR employs a specially trained, protein-protein interaction-sensitive large protein language model to extract intra-chain and inter-chain residue-residue contact information and evolutionary relationships, eliminating the need for laborious MSA searches. It also incorporates innovatively designed structure prediction and representation-driven flexible docking modules to enhance structure prediction, particularly for interacting contacts. tFold-TCR can predict unliganded TCR complexes, unbound pMHC complexes, and TCR-pMHC complexes within seconds. To provide insights into one of the most diverse receptor-ligand interactions in biology, we leverage tFold-TCR’s high-throughput prediction capabilities to construct TCRStructDB, the largest database providing TCR-pMHC structures, encompassing almost all currently known paired TCR-pMHCs and paired unliganded TCRs and pMHCs. We discuss the reliability of the TCRStructDB data characteristics and provide an online platform to expand its applications, supporting T cell-mediated cancer treatment and improving the success rate of immunotherapeutic interventions. Last but not least, we also explore a potential approach for designing the CDRs of TCRs using tFold-TCR.

## 2 Results

In this section, we first present the performance of tFold-TCR in predicting the structures of TCRs and pMHC in Section 2.1. Subsequently, we discuss the performance of tFold-TCR in predicting the structures of TCR-pMHC complexes in Section 2.2. Using the high accuracy and computational efficiency of tFold-TCR in predicting TCR-related structural systems, we compiled sequence pair information for TCRs, pMHCs, and TCR-pMHC complexes. This effort culminated in the creation of the largest TCR structural dataset, termed TCRStructDB, the details of which are elaborated in Section 2.3. In Section 2.4, we examine the characteristics of the TCRStructDB data, which includes both sequence and structural data, and outline the various applications of TCRStructDB.

### 2.1 Sequence-based structure prediction for unliganded TCR and unbound pMHC

Due to the limited of TCR-system structural data in the Protein Data Bank (PDB), tFold-TCR employs separate modules to predict TCR and pMHC structures independently, rather than using a single shared model for different inputs as methods like AlphaFold-Multimer and AlphaFold-3 do. Accurately predicting the structures of TCR and pMHC, particularly the CDR3 region of TCR, is crucial for validating their functions and is a prerequisite for tFold-TCR to accurately model the TCR-pMHC complex structures.

In tFold-TCR, the models for predicting unliganded TCR and unbound pMHC share the same architecture and are initialized with identical parameters. The primary distinction between these models lies in the use of different training datasets. Taking the unliganded TCR prediction model as an example, as illustrated in Fig. 1a, it comprises three main modules: a pre-trained large protein language model (ESM-PPI-TCR), a feature refinement module (Evoformer-Single) and a structure module. The ESM-PPI-TCR module is designed to extract both intra-chain and inter-chain information from the protein complex, generating features for downstream structure prediction tasks. We developed ESM-PPI-TCR by extending the existing ESM-2 model [25] through further pre-training using both monomers and multimers curated from several large databases, including UniRef50 [36], PDB, the Protein-Protein Interaction (PPI) database, the Antibody database, the peptide database, and our collected TCR database (refer to the Appendix C.2 for details). This enhancement enables the model to effectively extract intra-chain and inter-chain information, as well as peptide inter-chain information for pMHC. The Evoformer-Single module updates and refines the input protein features derived from the language model. Finally, the structure module employs invariant point attention (IPA) [17] to perform SE(3) equivariant transformations, mapping the obtained representations to the predicted atomic-level 3D structure. This modular approach ensures that tFold-TCR can accurately predict the structures of unliganded TCR and unbound pMHC, which is crucial for the subsequent accurate modeling of TCR-pMHC complex structures (Details in the Section 2.2).

**Fig. 1:**
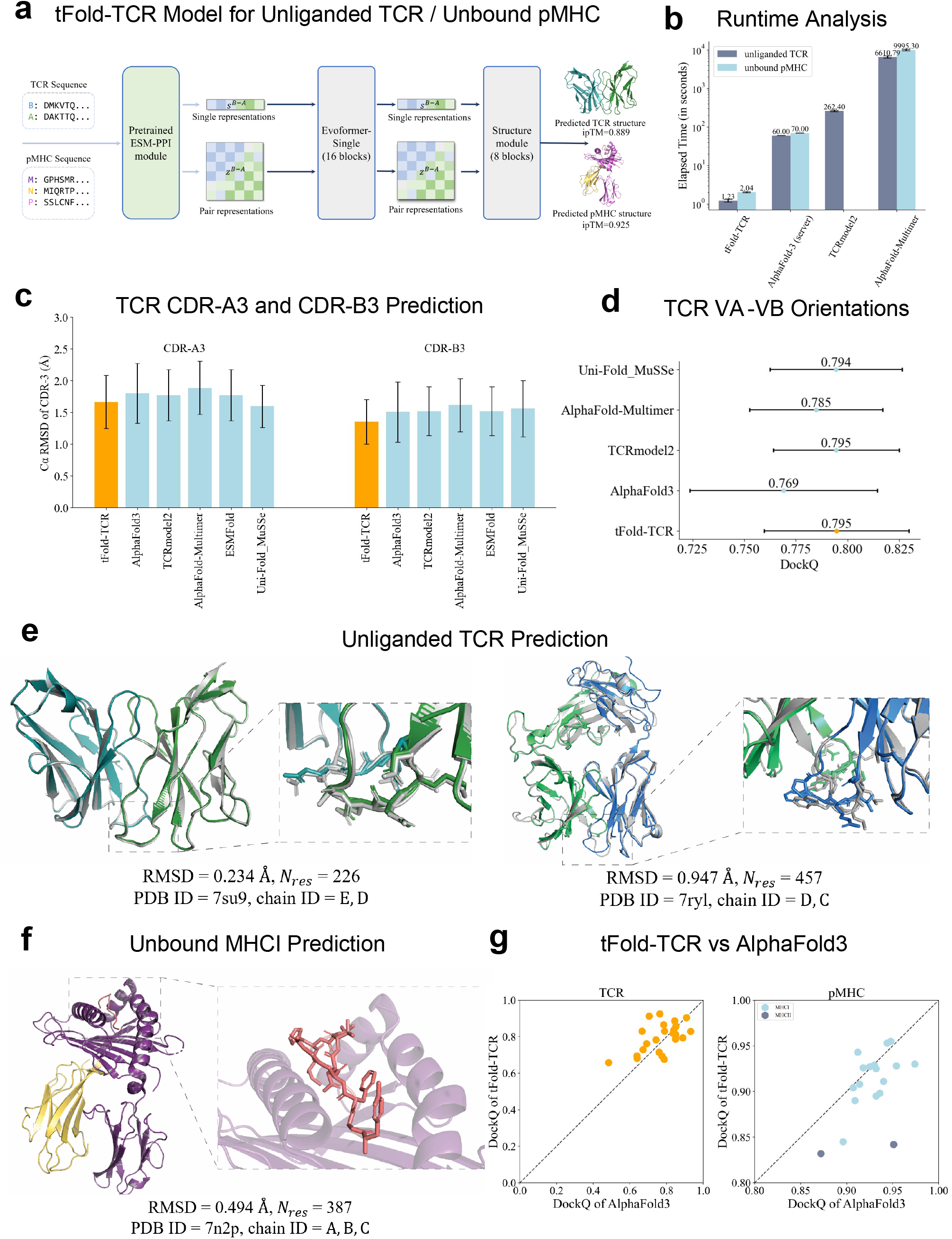
tFold-TCR method and result for unliganded TCR and unbound pMHC. (a) Overview of the tFold-TCR architecture for unliganded TCR and unbound pMHC, with arrows indicating the direction of information flow. Gradient backpropagation is only enabled for dark arrows, but not for light arrows. (b) Runtime analysis for tFold-TCR on 24 TCRs in STCRDab-22-TCR and 18 pMHCs in STCRDab-22-pMHC. Comparison to AlphaFold-3, TCRModel2 and AlphaFold-Multimer. All runtime are measured on a single NVIDIA A100 GPU with 20 CPU cores except AlphaFold-3, which is run on AlphaFold Server. (c) Performance comparison of tFold-TCR with other structure prediction method on STCRDab-22-TCR test set, evaluated by backbone RMSD in CDR-A3 and CDR-B3, represented as mean data with a 95% confidence interval. (d) DockQ evaluation performance on the SAbDab-22H2-TCR. (e) Comparison of our predicted structures for unliganded TCR targets (PDB 7su9 and 7ryl, blue for the beta/gamma chain, green for alpha/delta chain) with their respective experimental structures (gray). The accurate prediction of the CDR region by tFold-TCR for both side and main chains is particularly noteworthy. (f) Comparison of our predicted structures for unbound pMHC targets (PDB 7n2p, purple and yellow for the MHC chains, red for the peptide chains with their respective experimental structures (gray). The accurate prediction of the flexible peptide by tFold-TCR is particularly noteworthy. (g) Head-to-head comparison between tFold-TCR and AlphaFold-3 on STCRDab-22-TCR and STCRDab-22-pMHC datasets.

To comprehensively evaluate the performance of tFold-TCR, we employed a widely adopted temporal separation method. Using a cutoff date of January 1, 2022, data prior to this date were used for training and validation, while data from 2022 onwards were reserved for testing. For unliganded TCR and unbound pMHC, we constructed two non-redundant benchmarks: STCRDab-22-TCR and STCRDab-22-pMHC, comprising 24 TCRs and 18 pMHCs, respectively.

We first compared the inference time of tFold-TCR with that of existing methods, as illustrated in Fig. 1b. Notably, tFold-TCR demonstrates a remarkable speed advantage, being 5000 times faster than AlphaFold-Multimer [18] for unliganded TCRs and 4000 times faster for unbound pMHCs. This significant improvement in speed can be attributed primarily to the implementation of ESM-PPI-TCR, which effectively eliminates the need for time-consuming multiple sequence alignment (MSA) searches. Although AlphaFold-3 also utilizes MSA, it has undergone extensive engineering optimizations that mitigate its speed disadvantage, resulting in tFold-TCR being approximately 40 times faster than AlphaFold-3.

For general protein structure prediction, PLM-based models are known for their speed but often exhibit slightly lower performance compared to MSA-based models. However, as illustrated in Fig. 1c and Table. B2, tFold-TCR demonstrates exceptional performance in predicting the structure of unliganded TCRs. Specifically, it achieves average root-mean-squared-deviation (RMSD) values of 0.566 Å and 0.469 Å for the framework regions (FR) in the alpha chains and beta chains, respectively. In the more difficult task of predicting the CDRs, tFold-TCR achieves average RMSD values of 0.736 Å, 0.560 Å, and 1.665 Å in the CDR-1, CDR-2, and CDR-3 regions of the alpha chains (denoted as A1, A2, and A3 regions), respectively. For the corresponding CDRs of the beta chains (denoted as B1, B2, and B3 regions), the model achieves RMSD values of 0.442 Å, 0.441 Å, and 1.352 Å, respectively. All RMSD scores are calculated over the backbone beta atoms, following the alignment of the respective framework residues. We compared tFold-TCR with currently existing general protein structure prediction methods, including AlphaFold-Multimer [18], AlphaFold-3 [19], as well as TCR-specific methods, including TCRModel2 [21]. In the FR, all examined methods consistently exhibited the highest performance compared to other regions, but tFold-TCR proved to be the most accurate among them. The CDR-A3 and CDR-B3 regions are the most challenging components for prediction due to their significant sequence and structural diversities. tFold-TCR significantly outperformed all other methods. This demonstrates tFold-TCR’s superior capability in handling the complex and diverse nature of these critical regions, further establishing its robustness and accuracy in TCR structure prediction. We also found that for longer CDR3 regions, tFold-TCR’s predicted CDR3-RMSD values increased, with a Pearson correlation coefficient of 0.7 (as shown in Fig. A1), which is consistent with our expectations, as longer CDR3 regions exhibit greater flexibility and are more challenging to predict.

The anticipated orientation between different paired TCR chains is crucial for determining the overall binding surface. To assess the accuracy of the inter-chain orientation, we used several metrics, including DockQ, the fraction of native contacts (Fnat), ligand root-mean-square deviation (LRMS), and interface root-mean-square deviation (iRMS), to evaluate the docking interface accuracy. Additionally, we used RMSD and GDT (Global Distance Test) to assess the overall structure of unliganded TCRs. tFold-TCR achieved the best performance among the evaluated methods in most metrics, except for Fnat. Specifically, tFold-TCR’s predicted overall structure is the most accurate, with an average RMSD of 0.704 Å and an average GDT of 0.956. These results highlight tFold-TCR’s superior capability in accurately predicting both the inter-chain orientation and the overall structure of unliganded TCRs, making it a highly reliable tool for TCR structure prediction.

While tFold-TCR demonstrates rapid and accurate predictions for unliganded TCRs, its performance for unbound pMHCs is slightly inferior to MSA-based methods such as AlphaFold-Multimer and AlphaFold-3. As shown in Table. B3, the average DockQ score for tFold-TCR on the STCRDab-22-pMHC dataset is 0.908, compared to 0.927 and 0.926 for AlphaFold-Multimer and AlphaFold-3, respectively. Although tFold-TCR achieves high overall structural accuracy (RMSD=0.684 Å, GDT=0.959), its prediction accuracy for peptide side chains is not as high as that of AlphaFold-3, resulting in a lower Fnat score. Despite the limited co-evolutionary information available for peptides, MSA-based models can extract more effective co-evolutionary signals through appropriate paired strategies, leading to more accurate predictions. Additionally, accurate template structures can help AlphaFold-Multimer and AlphaFold-3 optimize the side chain of peptide.

Fig. 1e provides examples of the predicted structures for *αβ*TCR and *γδ*TCR, offering a visual representation of the prediction results. The experimental results demonstrate that tFold-TCR can deliver high-precision predictions for CDR- A3, CDR-B3, CDR-D3, and CDR-G3 regions. Both the backbone and side chains of these regions show a high degree of similarity to the actual structures, which is particularly noteworthy given that these are the most challenging areas in TCR structure prediction. Fig. 1f illustrate examples of predicted unbound pMHC, along with the corresponding experimental structures, demonstrating that tFold-TCR can provide high-quality predictions of the interaction interfaces between peptide and MHCs.

Fig. 1g presents a head-to-head comparison between tFold-TCR and the state-of-the-art structure prediction method AlphaFold-3. The results indicate that tFold-TCR performs slightly better than AlphaFold-3 on unliganded TCRs, while it is marginally less accurate on unbound pMHCs. Notably, tFold-TCR’s prediction accuracy for MHCII-type pMHC structures is particularly lower compared to AlphaFold-3. The reduced performance can be attributed to the distinct structural and binding characteristics of MHC-I and MHC-II molecules, as discussed in Section 3.

### 2.2 High-throughput TCR-pMHC complex prediction using tFold-TCR

Once we trained a TCR structure prediction model and a pMHC structure prediction model, we used the outputs of these models as feature generation modules and integrated them using an additional flexible docking model. The full architecture of tFold-TCR, the constructed TCR-pMHC complex prediction model, is depicted in Fig. 2a. It mainly consists of three modules: the TCR feature generation module, the pMHC feature generation module, and the representation-driven flexible docking module.

**Fig. 2:**
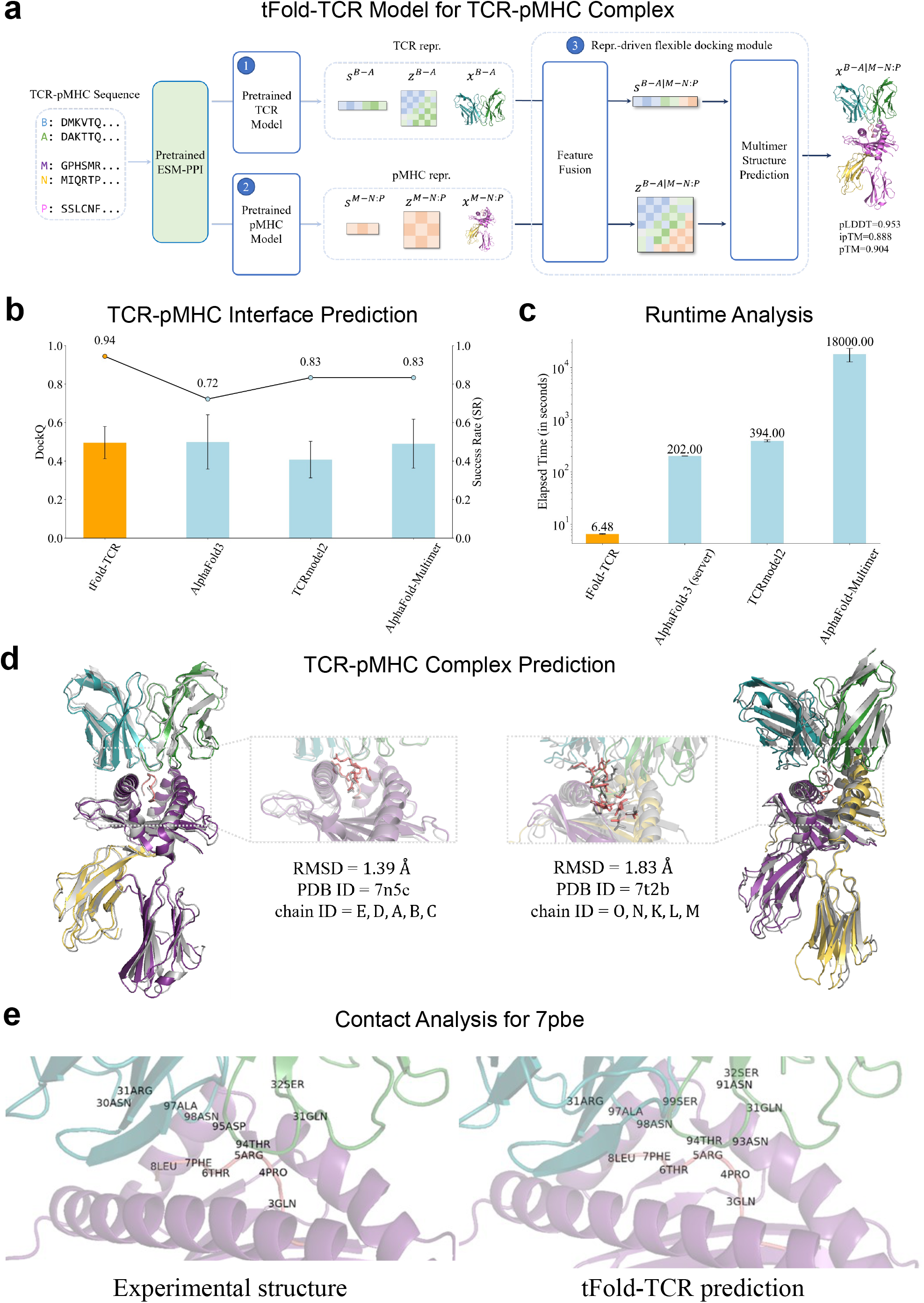
tFold-TCR method and result for TCR-pMHC complex. (a) Overview of the tFold-TCR architecture for TCR-pMHC complex, with arrows indicating the direction of information flow. Gradient backpropagation is only enabled for dark arrows, but not for light arrows (Repr. : Representation). (b) Performance comparison of tFold-TCR with other structure prediction methods on STCRDab-22-TCR pMHC test set, evaluated by DockQ and success rate. (c) Runtime analysis for tFold-TCR on 18 TCR-pMHC complexes in STCRDab-22-TCR pMHC. Comparison to AlphaFold-Multimer, AlphaFold-3, TCRModel2. All runtime are measured on a single NVIDIA A100 GPU with 20 CPU cores except AlphaFold-3, which is run on AlphaFold Server. (d) Comparison of our predicted structures for TCR-pMHC complex targets (PDB 7n5c and 7t2b, blue for the beta chain, green for alpha chain, purple and yellow for the MHC chains and red for the peptide chains) with their respective experimental structures (gray). The accurate prediction of the interface between TCR and pMHC by tFold-TCR for both side and main chains is particularly noteworthy. (e) An example of a tFold-TCR prediction structure (left) and experimental structure (right) for TCR-pMHC complex targets (PDB ID: 7pbe, blue for the beta chain, green for alpha chain, purple for the MHC chain and red for the peptide chains), with binding contacts between TCRs and peptide are highlighted.

During the training of tFold-TCR for complex prediction, the pre-trained feature generation modules (including both TCR and pMHC) remain fixed to facilitate the convergence and optimization of the entire model. The representation-driven flexible docking module is primarily composed of two parts: the feature fusion module and the complex structure prediction module. The feature fusion module effectively integrates the sequence representation, pair representation, and the unbound structures of TCR and pMHC obtained from the TCR and pMHC feature generation modules. This integration generates the initial sequence representation and pair representation of the TCR-pMHC complex. Subsequently, the complex structure prediction module, which includes an Evoformer-Single stack with 16 blocks and a structure module with 8 blocks, updates the representations via the Evoformer-Single blocks. It then maps these updated representations to the predicted complex structure and provides the predicted confidences via the structure blocks. In the representation-driven flexible docking module, tFold-TCR not only computes the conformation of the TCR-pMHC complex but also updates the structures of TCR and pMHC, simulating the process of flexible docking. Importantly, tFold-TCR does not require the input TCR and pMHC to be paired, as it utilizes positional encoding for different chains, allowing it to effectively identify and accommodate missing chains (e.g., TCRs with only the Beta chain or MHCI with a single chain). This capability allows it to efficiently utilize structural data from the STCRDab [37] database.

Similar to the data preparation process in unliganded TCR structure prediction, we curated a hold-out test set to evaluate the performance of structure prediction methods on TCR-pMHC complexes from the STCRDab [37], employing a temporally separated approach. To ensure a fair comparison with existing methods such as AlphaFold-Multimer and AlphaFold-3, we made sure that none of the evaluated methods had been exposed to the structures included in the test set by setting the cutoff date from January 1, 2022, to December 31, 2022. The finalized test set, named STCRDab- 22-TCR pMHC, includes 18 TCR-pMHC complexes. Considering the relatively limited structural data available in STCRDab, we additionally constructed an in-house benchmark. This benchmark, named Rossjohn Lab Benchmark, contains 6 unpublished TCR-pMHC complexes, and all structures are not present in databases such as PDB or STCRDab. This dataset is more challenging compared to STCRDab-22-TCR pMHC.

We compared tFold-TCR with the famous structure prediction methods, including AlphaFold-Multimer [18] and AlphaFold-3 [19], as well as TCR-specific methods like TCRModel2 [21] and TCRDock [20]. Additionally, we evaluated several docking methods [38–41]. However, even when using the bound TCR and bound pMHC as inputs, the performances of these docking methods were significantly inferior.

As shown in Fig. 2b, Table. B4 and Table. B5, tFold-TCR performs comparably to AlphaFold-3 and AlphaFold-Multimer in predicting TCR-pMHC complex structures. On the STCRDab-22-TCR pMHC benchmark, tFold-TCR achieved a DockQ score of 0.494, while AlphaFold-3 and AlphaFold-Multimer scored 0.490 and 0.494, respectively. Similar conclusions were drawn from the Rossjohbn Lab Benchmark, where the average DockQ scores for tFold-TCR, AlphaFold-3, and AlphaFold-Multimer were 0.474, 0.470, and 0.439, respectively. Despite all three methods assigning high confidence scores (evaluated by ipTM, confidence score) above 0.9 to their predicted structures, tFold-TCR demonstrated more robust performance, with a success rate (SR) approaching 100%. tFold-TCR maintains a stable prediction accuracy compared to AlphaFold-3 and AlphaFold-Multimer, without hallucination behavior described in [19]. Head-to-head comparison between tFold-TCR and the AlphaFold-3 and AlphaFold-Multimer are presented in Fig A2. Compared to TCR-MHCI, the accuracy of tFold-TCR in predicting TCR-MHCII is relatively lower. This discrepancy can be attributed to the open structure of MHC-II molecules, which allows the peptide ligand to extend from both ends of the binding groove. This characteristic increases the complexity of the TCR-pMHC-II binding motif, making accurate predictions more challenging. For TCRModel2, the pMHC is truncated, retaining only the two alpha helices and one beta sheet structure at the top of the MHC. TCR-specific MSA and templates are used as inputs for AlphaFold-Multimer to predict the structure. This truncation of the pMHC makes the comparison somewhat unfair. When evaluated using the complete TCR-pMHC complex, the average DockQ score is approximately 0.4.

The performance advantage of tFold-TCR compared to AlphaFold-Multimer and AlphaFold-3 is not particularly significant. However, the greatest advantage of tFold-TCR lies in its computational speed. Since tFold-TCR eliminates the need for time-consuming MSA searches and instead uses a language model as input, tFold-TCR is over 2000 times faster than AlphaFold-Multimer and 30 times faster than AlphaFold-3, as shown in Fig 2c. The ability to quickly and accurately predict structures allows tFold-TCR to be widely applied to various tasks, one of which will be introduced in Section 2.3. We also used tFold-TCR to predict the structures of unliganded TCRs in the STCRDab-22-TCR pMHC dataset and com- pared the results with those of the TCR pMHC complex. We observed that employing the representation-driven flexible docking module for joint modeling of the TCR-pMHC system resulted in a 4% reduction in the RMSD of the TCR CDR- A3 and CDR-B3 regions. Although this improvement may not seem substantial, it highlights the role of flexibility in enhancing overall structural modeling.

In the structural prediction of TCR-pMHC complexes, accurately predicting the interaction contacts between CDR loops and epitopes is of paramount importance, yet it remains a challenging task due to the disordered and flexible nature of these regions. Fig. 2d illustrates examples of the predicted structures for TCR-MHCI and TCR-MHCII, highlighting the contact regions. Notably, while tFold-TCR aligns with most structural prediction tools, it shares similar limitations in accurately predicting side-chain interactions [42]. However, the backbone interface between TCR and pMHC is accurately predicted by tFold-TCR. For the example shown in Fig.2e, the experimental structure reveals a total of 14 contact pairs, while tFold predicts 15 contact pairs. The performance metrics indicate a recall of 12 out of 14, resulting in a recall rate of approximately 85.7%, and a precision of 12 out of 15, yielding a precision rate of 80%. The DockQ score is 0.76, and the iRMS is 1.16. Importantly, tFold-TCR’s predictions effectively reproduce the CDR loop interactions with the P4-P6 epitope residues, which is critical for the specific recognition of peptides. Understanding these interactions is essential for identifying potential viral escape mutations from common public TCRs.

### 2.3 Large-scale structural database of TCR and pMHC

The protein structure is indicative of its function and activity [12]. To elucidate the structural characterization of TCR-pMHC interactions, we developed TCRStructDB, the largest structural TCR database available to date, based on the newly developed tFold-TCR. TCRStructDB is an online platform offering extensive data, including structures of known TCR-pMHC complexes, paired unliganded TCRs and unbound pMHCs, and a suite of search and analytical tools.

We collected paired full-chain TCR-pMHC complexes, unliganded TCRs, and pMHC sequencing data from existing databases such as STCRDab [37], TCR3d [43], IEDB [44], HuARdb [45], OTS [46], as well as various studies utilizing 10x sequencing to profile immune cells. The data availability information provided by the original publications guided our collection process (see Section 4.5 for details). We manually curated metadata from these publications with well-established data-processing pipelines. When experimental structures were not available, we used tFold-TCR predictions to provide the corresponding structures for the included sequencing data. Specifically, during the database curation process, we applied a uniform processing pipeline to all raw data, as illustrated in Fig. 3a. The pipeline consists of three main stages: raw sequencing data processing, annotation and validation of sequence data, and structure prediction.

**Fig. 3:**
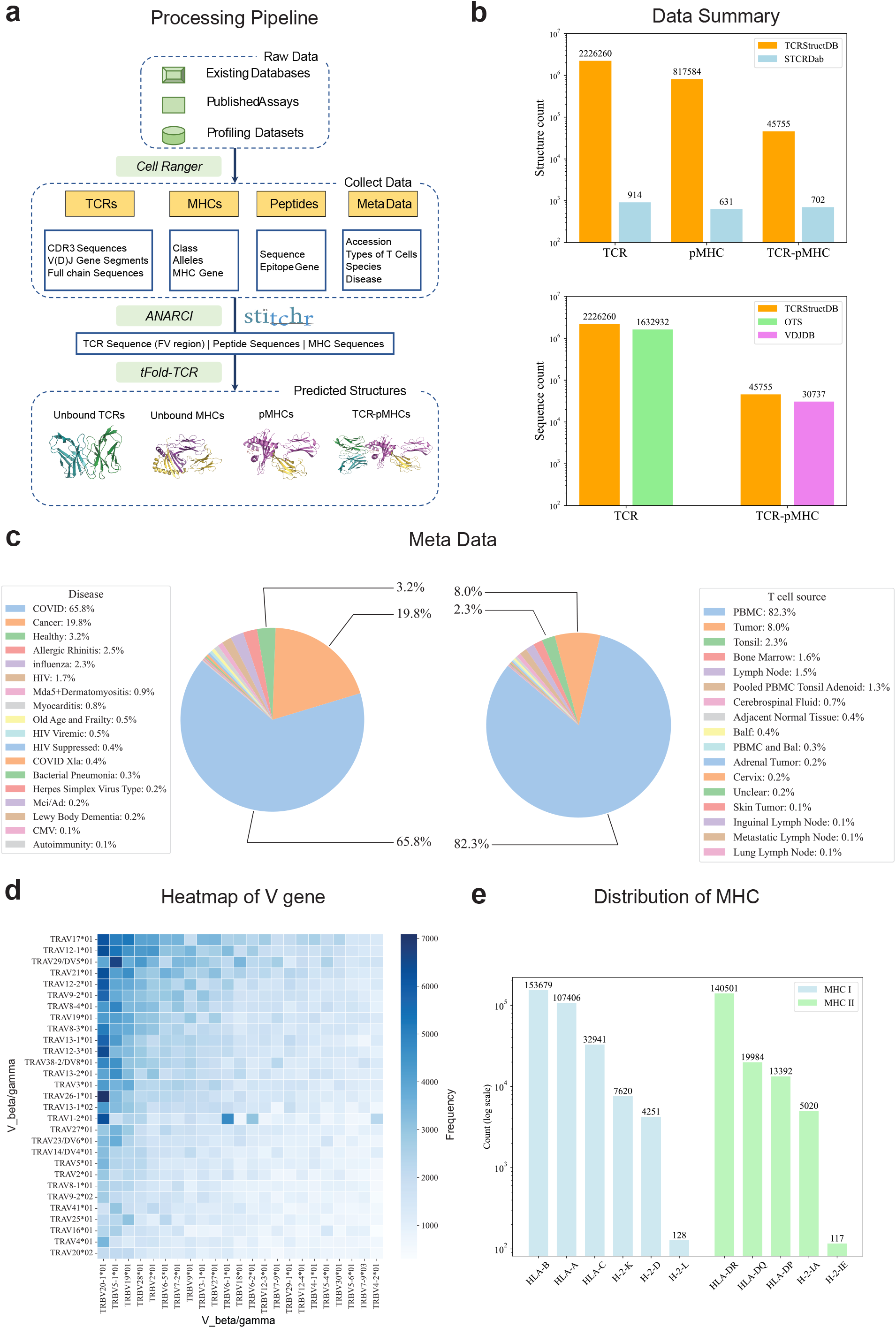
Overview of TCRStructDB. (a) Data collection and processing pipeline for TCRStructDB. The pipeline integrates data from existing databases, published assays, and profiling datasets, utilizing tools such as Cell Ranger, Stitchr, and ANARCI to generate TCR, peptide and MHC sequences. It employs tFold-TCR to generate 3D structures of unliganded TCR, unbound MHC, pMHC, and TCR-pMHC complexes. (b) Comparison of data amount between TCRStructDB and existing databases, including structural databases (top) and sequence databases (bottom). TCRStructDB exhibits a sub- stantial increase in the number of entries for TCR, pMHC, and TCR-pMHC compared to STCRDab, OTS, and VDJDB. (c) Metadata analysis of TCRStructDB. Pie charts illustrate the distribution of diseases and T cell sources within the database. Categories representing less than 0.1% are not shown. The top three categories are labeled with their respective proportions. (d) Heatmap of V gene pairing in TCR alpha and beta chains, displaying the top 75% of gene pairings based on frequency in the database. (e) Distribution of MHC class I and II subtypes in the pMHC data.

In the first stage, we processed the raw sequencing data to obtain the TCR-relevant amino acid (AA) sequence data. Specifically, the downloaded raw 10X sequencing data were re-analyzed using the 10x Genomics CellRanger software. Individual sequences were then paired by referring to the 10x barcodes. In the second stage, software such as ANARCI [47] was utilized to renumber TCR sequences, ensuring that they were full-length and free of ambiguous amino acids. For cases where full-chain sequences of TCRs could not be obtained, Stitchr [48] was employed to generate valid full-chain TCR sequences based on the available V and J gene data, CDR3 sequence information, and species meta- data. Following this, ANARCI-based annotation and validation were performed. In the final stage, we utilized tFold-TCR to obtain the structural information for various TCR-relevant sequences, including unliganded TCRs, unbound MHCs, pMHCs, and TCR-pMHC complexes (refer to Section 4.5 for details).

As a result, TCRStructDB currently contains a total of 4,517,910 paired TCR sequences, including 2,226,260 unique TCR fragment variable (Fv) region sequences, each matched with its corresponding predicted structural data. Additionally, we have predicted structural data for 817,584 pMHC complexes, comprising 531,578 entries for MHC I and 286,006 for MHC II, as well as 45,755 predicted structural data entries for TCR-pMHC complexes. Compared to the current existing TCR-specific structural database such as STCRDab [37], TCRStructDB provides significantly more structural data, offering an increase of approximately 2400-fold of TCR, 1300-fold of pMHC and 65-fold of TCR-pMHC data compared to STCRDab. In comparison to the largest known TCR-specific sequence databases, OTS [46] and VDJDB [49], TCRStructDB encompasses all their data, thereby surpassing them in richness and comprehensiveness (Fig. 3b). Furthermore, TCRStructDB will be continually updated and maintained to increase the depth and diversity of available data.

Analyzing the data in TCRStructDB, we found that the currently available data exhibit a relatively wide distribution of diseases and T cell sources (Fig. 3c). COVID-19 and cancer cases are the two most studied sample sources, making up 65.8% and 19.8% of all samples, respectively. Additionally, among these sources, the majority of T cell data are derived from peripheral blood mononuclear cells (PBMCs), accounting for approximately 82.3%, followed by tumor tissues at around 8%. In addition, the distribution of T cell type and T cell subtype are shown in Fig. A3.

The Fv region of the TCR is generated by the rearrangement of variable (V) genes and joining (J) genes. When analyzing the V-allele pairing in non-redundant sequences, we observed dramatic frequency differences among different V-allele pairs. Specifically, V-allele pairs such as TRAV29/DV5*01—TRBV5-1*01, TRAV1-2*01—TRBV20-1*01, TRAV17*01—TRBV20-1*01 were found to be among the top-ranked pairs (Fig. 3d). This may be attributed to the combination of these genotypes may be more likely to occur during TCR gene rearrangement. In terms of MHC subtypes, HLA-B was identified as the most prevalent MHC-I subtype, while HLA-DR was the most frequent MHC-II subtype (Fig. 3e).

### 2.4 Data characteristics and applications of TCRStructDB

TCRStructDB, to our knowledge, is the largest database to date, offering paired sequence and structural information for TCR-pMHCs, TCRs and pMHCs, which has the potential to provide valuable insights through data analysis, similar to previous works [50–52] based on AlphaFoldDB [31] and ESM Atlas [25]. We first analyzed the functional characteristics of TCRs from a sequence perspective. Our analysis revealed that TCRs sharing the same V gene and J gene were clustered together in the sequence-feature space derived from sequence-based language models (ESM-PPI-TCR). However, we did not observe clustering of TCRs that bind to the same pMHC, indicating that sequence similarity alone may not fully account for the binding specificity of TCRs to their respective pMHCs (refer to supplementary section C.2.3 for details). This suggests that additional factors, such as structural features or the dynamics of TCR-pMHC interactions, may play a significant role in determining TCR specificity and functionality. In this section, we further analyzed the functional characteristics of TCRs from a structural perspective to explore biological insights and potential applications provided by TCRStructDB.

The docking angles between TCR and pMHC, including the crossing angle and incident angle, are key geometric parameters utilized in the analysis of TCR-pMHC complex [53]. Despite the variability in the formation of TCR-pMHC complexes, existing structural studies [54–56] demonstrate that TCRs maintain a consistent docking orientation in complex with different MHC molecules falling within a tight distribution of angles, clustered across different structures. Previous studies [57] suggest that TCRs selected during thymic development are restricted to recognizing pMHC complexes in a specific docking orientation, which is essential for normal immune function. Most TCRs that recognize class I or class II MHCs focus their binding interactions on the central regions of the *α*1- and *α*2/*β*1-helices. However, an existing study [57] found no skewed usage of TCR V genes within MHC allele groups and no conserved TCR-MHC contacts in MHCI. This finding challenges the hypothesis that TCR and MHC genes co-evolved for germline recognition. In fact, it suggested that the docking orientation is largely influenced by conserved residues in specific regions of MHC molecules [58]. It remains unclear whether the docking orientation preference is directly determined by germline-encoded residues or is a consequence of thymic selection [53].

To address this question, we conducted a docking angle analysis of TCR-pMHC complexes using both STCRDab, which contains a relatively limited number of experimental structures, and TCRStructDB, which offers a significantly larger set of predicted structures. We analyzed the correlations between the docking angles and the TRAV gene, TRBV gene, and MHC type. Analysis of variance (ANOVA) results illustrated in Fig. 4 and Table B7, show that the docking angles, including the crossing angle and incident angle, are highly associated with both the combination of TRAV and TRBV genes and MHC type significantly. The docking angle analysis of experimental structures in STCRDab yielded similar conclusions; however, we did not observe a significant association between the incident angle and MHC type (Table B6). This lack of association may be attributed to the limited sample size in the experimental database. Additionally, we modeled the distribution of docking angles, as illustrated in Fig. A4. Our findings indicate that the docking angle distribution can be accurately fitted using both the V and D genes, as well as MHC type, but not by the MHC type alone. Based on these findings, we hypothesize that the docking angles are determined by thymic selection, where TCRs with suitable docking angles for the given MHC type are retained. Specific regions of the MHC are crucial in determining the classic TCR-pMHC docking orientation. Different TRAV-TRBV paired TCR molecules then form unique binding patterns on the MHC surface, defining the fine-tuned docking angles. Additional factors, such as interactions with coreceptors and electrostatic steering, may also play a role.

**Fig. 4:**
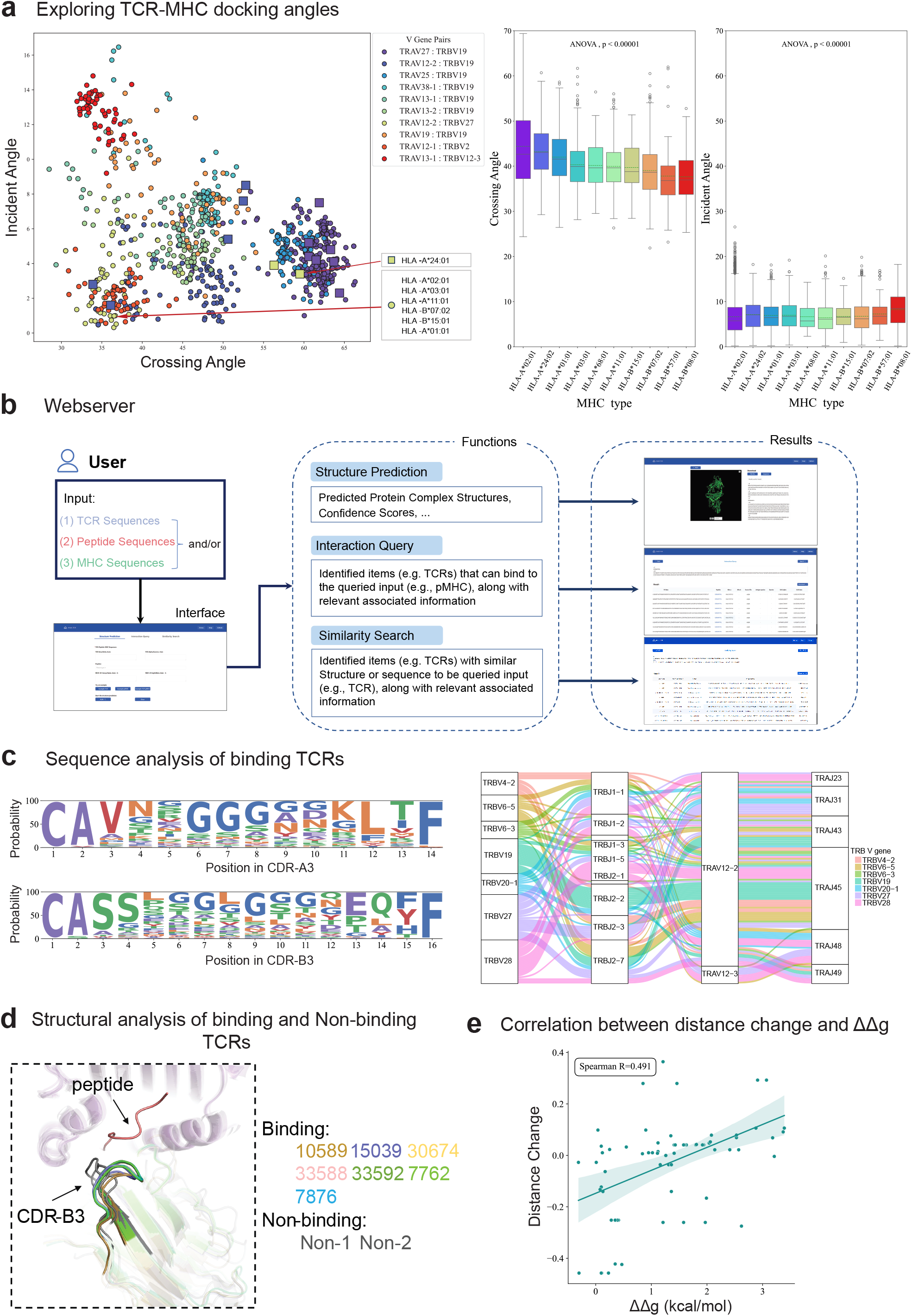
Data characteristics and applications of TCRStructDB. (a) Docking angle analysis. Left: Calculated docking angles (incident angle, crossing angle) for TCR-pMHC structures in TCRStructDB (prototype) and STCRDab (square), classified by TCR V gene pairs. The top 10 V gene pairs are displayed. TCRStructDB and STCRDab exhibit similar data distributions, with one outlier MHC type shown, indicating that MHC, in addition to TCR, also influences the docking angle. Right: ANOVA analysis of the correlation between docking angles of TCR-pMHC structures in TCRStructDB and MHC type. (b) Schematic overview of the TCRStructDB webserver. The server encompasses three main functionalities: structure prediction, binding retrieval, and similarity retrieval. Users can search by TCR sequence or pMHC sequence/alleles, and the server provides structural results, sequence metadata, and similar sequence analysis. (c) Sequence analysis of binding TCRs of MART-1 protein. Left: Distribution of CDR-A3 (top) and CDR-B3 (bottom) amino acid types for binding TCRs. Right: The visualized figure of V-J genes pairing relationship for binding TCRs. (d) Structure analysis of binding and non-binding TCRs of MART-1 protein. Binding and non-binding TCRs have different docking angles. (e) Correlation between distance change and affinity changes. The bands represent the 95% confidence intervals for the linear regression fit, showing the correlations between the average distance changes of residue pairs predicted by tFold-TCR and the experimentally determined affinity changes (ΔΔ*G*) of mutant samples.

To facilitate the utilization of TCRStructDB, we developed it as an online platform that offers a suite of search and analytical web applications. These applications are categorized into three main functional groups: structure prediction, binding retrieval, and similarity retrieval. As illustrated in Fig. 4b, users input their query TCR and pMHC sequences into the interface. The corresponding function is then invoked through the appropriate interface, and after computation, the backend displays the relevant results, including predicted structures and database retrieval outcomes. Detailed information for the TCRStructDB online platform is introduced in the Supplementary C.4.

ELAGIGILTV (MART-1 protein from melanoma) is a crucial tumor-associated antigen presented to T cells by MHC and has been extensively studied [59]. Using the MART-1 target, we searched the TCRStructDB database and retrieved a series of TCRs capable of binding to this target. Fig 4c analyzes these TCR sequences. In the CDR-A3 region, sequences predominantly start with “CAV” and end with “LTF”, with a high frequency of “GGG” in the middle positions, suggesting a conserved motif critical for binding specificity. In the CDR-B3 region, sequences predominantly start with “CASS” and end with “EQYF”, with notable conservation of “GGG” and “G” at various positions, indicating a recurring pattern essential for interaction with the MART-1 antigen. For V and J gene usage, the TRAV12-2 gene is dominant in the A chain, while no dominant V gene is observed in the B chain, with TRBV27 and TRBV28 showing higher frequencies.

In addition to sequence data, the unique feature of TCRStructDB is its ability to analyze the structures of TCRs that can bind to antigens, providing researchers with an additional perspective. We further collected a subset of TCRs that cannot bind to this target from 10x Antigen-Peptide Specific Stimulation Experiments [60] and predicted the corresponding TCR-pMHC structures. These were compared with the TCR-pMHC structures from TCRStructDB that can bind to the target, and a random selection of sequences was visualized, as shown in Fig 4d. We found that TCRs capable of binding to the MART-1 antigen exhibit similar CDR-B3 structures, which differ significantly from the structures of non-binding TCRs, displaying completely different docking orientations.

The combined sequence and structure analysis of binding and non-binding TCRs is significant as it provides comprehensive insights into the determinants of antigen recognition and binding. By identifying conserved motifs and structural patterns, we can better understand the specificity and affinity of TCR interactions with tumor-associated antigens.

In addition to structure, affinity has always been a key concern for researchers. However, the affinity between TCR and pMHC is generally weak, and known affinity data is scarce, which is not included in TCRStructDB. Similar to most structural prediction models, our model predicts a binding structure regardless of whether the input TCR can bind to the pMHC or whether the peptide can bind to the MHC. However, given that our model can achieve complex predictions in seconds, we can provide additional input information to those sequence-based binding prediction models and affinity prediction models [61, 62].

To validate our conclusions, we obtained 76 mutant sequences and their corresponding mutation effects based on the 1AO7 [54] complex from the ATLAS [63] database, as 1AO7 has the most mutations available in the database. Most of these complexes do not have accurate experimental structures. Given that TCR specificity primarily arises from CDR3- peptide interactions, only mutations occurring in the CDR3 and peptide regions were retained. Using tFold-TCR, we predicted the structures of these mutants and calculated the mean distances between the CDR3 segments and peptides. We analyzed the relationship between the predicted distance changes and the experimentally observed mutation effects (ΔΔ*G*). Our results revealed a moderate positive correlation (Spearman = 0.491), as shown in Fig 4e, indicating that tFold-TCR has potential for predicting changes in binding affinity resulting from protein mutations.

The accurate sequence and structural characteristics in TCRStructDB, combined with the applications of our online platform, potentially lead to more effective T cell-mediated cancer treatments. Additionally, the ability to distinguish between binding and non-binding TCRs at both the sequence and structural levels can enhance the precision of therapeutic TCR selection, improving the overall success rate of immunotherapeutic interventions.

## 3 Conclusion and discussion

In conclusion, tFold-TCR not only serves as an efficient tool for rapid and accurate structure prediction but also demonstrates potential as a platform technology that can drive innovation in TCR-based therapeutics and vaccines. By enabling the exploration of the extensive landscape of potential TCR-pMHC interactions, tFold-TCR facilitates the design of effective TCR-based treatments. Its ability to enhance the accuracy and speed of TCR-pMHC complex structure predictions can be leveraged for applications such as virtual screening and TCR design. We anticipate that tFold-TCR will provide immunologists with a comprehensive tool to accelerate advancements in structural bioinformatics. Additionally, the TCRStructDB constructed in this work, which is the largest TCR structure database to date, offers structural biologists a valuable data resource, enriching their understanding of these complex interactions. This resource has the potential to lead to significant insights in molecular biology and immunology, fostering breakthroughs in therapeutic applications.

In the following sections, we will explore a potential approach for designing the CDRs of TCRs using tFold-TCR, as well as discuss the limitations of our current model and outline future works.

### 3.1 TCR CDRs design using tFold-TCR

AI has been extensively utilized in antibody design [12, 64, 65], yet there are limited successful instances of AI-driven TCR design. Some approaches [66, 67] have employed language models for TCR design. Recent studies [29, 68] have shown that the co-design of the structure and sequence of antibodies can predict sequences that are close to real antibodies. We have extended our tFold-TCR model to design the CDR sequences of TCRs while predicting their structures.

In the design phase, masked regions indicate positions where new amino acids are to be generated. These regions are input into ESM-PPI-TCR as *<*Mask*>* tokens, resulting in a TCR representation with masked regions and an initial TCR structure where the masked regions contain only backbone atoms. Subsequently, tFold-TCR integrates the features of the masked TCR and the unmasked pMHC in the feature fusion module. Following updates in the Evoformer-Single Stack of the TCR-pMHC embedding, tFold-TCR employs an additional auxiliary head (implemented as feed-forward layers) to recover the amino acid types of the masked TCR regions. This architecture considers the interactions between TCR and pMHC, aiming to design TCRs that interact effectively with pMHC. For the TCR design task, we introduced Sequence Recovery Loss during the training phase and adopted a multi-task learning approach to simultaneously predict both sequence and structure. We also constructed a test set, and on this test set, the amino-acid recovery (AAR) for CDR- B3 and CDR-A3 using tFold-TCR was approximately 33% and 40%. This AAR is relatively low compared to antibodies, primarily due to the limited structural data available for TCRs compared to antibodies. This scarcity of data highlights the importance of our efforts to build a comprehensive TCR database. Detailed methods and experimental specifics regarding the TCR design are provided in the Appendix C.3.

### 3.2 Limitations and future works

Although tFold-TCR achieves high-precision modeling of the TCR-pMHC system in seconds and can be used to design CDRs, it still has several limitations.

Firstly, tFold-TCR’s prediction accuracy for MHC-II-type pMHC and TCR-pMHC structures is notably lower than that of AlphaFold-3. This discrepancy arises from two main factors. From a structural perspective, MHC-I typically presents shorter peptides with a clear binding motif, primarily due to the significant roles of pockets B and F, which preferentially bind anchoring residues at the second position (P2) and the C-terminal (PΩ). This simplifies the prediction of MHC-I pMHC structures. In contrast, MHC-II molecules bind longer peptides that fit more snugly in the groove, involving four pockets that interact with residues at positions P1, P4, P6, and P9 within the 9-amino-acid binding core [69]. This complexity complicates the accurate modeling of the MHC-II peptide binding motif [70]. Additionally, from a data perspective, AlphaFold-3 benefits from a larger training dataset, enhancing its generalization capabilities. In contrast, tFold-TCR’s limited exposure to MHC-II structures restricts its ability to accurately capture these complexities, as only 22% of the pMHC structures in the training set are MHC-II-type.

Secondly, the ipTM score is often used to evaluate the accuracy of docking predictions in structural modeling, and it has a strong correlation with DockQ values[18, 19]. However, for the TCR-pMHC system, confidence scores such as ipTM or pTM become less meaningful. In the case of tFold-TCR predictions, the ipTM score typically exceeds 0.8. When evaluating the STCRDab-22-TCR pMHC test set, the *R*^2^ value between the predicted DockQ and ipTM scores generated by tFold-TCR is only 0.11, compared to 0.06 for AlphaFold-Multimer. This discrepancy stems from the similar docking paradigms of TCR-pMHC interactions, and the weak correlation of these confidence scores limits our ability to utilize the predicted structures for self-distillation. Although tFold-TCR achieves nearly a 100% docking success rate, the proportion of high-quality predictions (DockQ *≥* 0.8) is relatively low. We also attempted to use actifpTM [71] to focus on the confident region of a TCR-pMHC interaction, but this did not improve the situation. Therefore, it is necessary to design unique confidence scores specifically for the TCR-pMHC system to better distinguish the predicted complex structures by tFold-TCR.

Additionally, the lack of metrics to determine whether a sequence pair can bind presents another challenge for our model. The affinity between TCRs and pMHCs is often low, and many TCRs can bind to pMHCs on the cell surface, triggering an immune response, but cannot be structurally resolved in vitro [72, 73]. As discussed in Section 2.4, tFold-TCR can perform structural predictions in seconds, thereby providing additional input information to sequence-based binding prediction models and affinity prediction models. We plan to develop related functionalities in the future.

## 4 Methods

### 4.1 Data

#### 4.1.1 STCRDab

We initially developed and evaluated the performance of our model using structural data from STCRDab [37], an automated and curated repository of TCR structural data derived from the PDB. During the training phase, we utilized 928 structures released prior to December 31, 2021, for both the training and validation sets, ensuring a fair comparison with AlphaFold-Multimer [18] and AlphaFold-3 [19]. we clustered the training and validation sets using MMseqs2 [74] based on 95% sequence identity, allocating 90% of the clusters for training and 10% for validation. We used ANARCI [47] with the IMGT [75] numbering scheme to trim the Fv region of the TCRs, while no trimming was performed for the pMHCs. During training and evaluation, we did not differentiate between TCR types or MHC types. Sequences containing non-standard amino acids and TCR sequences that could not be recognized by ANARCI were removed.

For the construction of the test set, we selected structures released by STCRDab between January 1, 2022, and December 31, 2022, to evaluate the performance of the trained model. We established three distinct test sets to assess our model’s predictive capabilities for unliganded TCRs, unbound pMHCs, and TCR-pMHC complexes, denoted as STCRDab-22- TCR, STCRDab-22-pMHC, and STCRDab-22-TCR pMHC, respectively. It is worth noting that some of the unbound structures were derived from bound structures, which may introduce a certain degree of evaluation bias. To prevent data leakage, we excluded all sequences from the test set that exhibited over 95% similarity to those in the training set, based on sequence identity calculated by MMseqs2 [74]. Furthermore, for each test set, we conducted clustering based on 95% sequence identity using the same tool, selecting one complex randomly from each cluster for inclusion in the test set. Ultimately, this process yielded 24 pairs of unliganded TCRs, 18 pairs of unbound pMHCs, and 18 pairs of TCR-pMHC complexes.

#### 4.1.2 Rossjohn Lab Benchmark

To further evaluate the performance of tFold-TCR in modelling TCR-pMHC complexes, we constructed an unpublished in-house benchmark derived from Rossjohn’s lab. This test set includes 6 TCR-pMHC complex structures, comprising five TCR-MHCI and one TCR-MHCII. None of the structures in this test set appear in the PDB. The sequence-to-sequence similarities between these structures and their closest counterparts in the PDB are 0.688, 0.752, 0.764, 0.812, and 0.845, as calculated by MMseqs2 [74]. These similarity scores highlight the challenge of this test set.

#### 4.1.3 General proteins in PDB for pre-training

The limited availability of structural data for TCRs constrains the model’s generalization capabilities. Recent studies [29, 76] have shown that pre-training on general protein monomers and multimers, followed by fine-tuning on antibodies, can improve the accuracy of antibody prediction. Our language model is trained on sequences of single or multiple chains, enabling it to extract features from both monomeric proteins and protein complexes. Consequently, it is justifiable to utilize complex structures from the PDB for the pre-training of our structural prediction model.

The PDB data construction methodology closely follows our previous work [29]. We collected all structures from the PDB released prior to January 1, 2022. Structures with a resolution exceeding 9Å or those with a proportion of missing residues greater than 0.8 were excluded from the dataset. We employed MMseqs2 [74] to cluster the remaining structures based on 40% sequence identity. The difference from our previous data construction approach is that, to better adapt the model to the pMHC system, we did not filter out peptides shorter than 40 amino acids.

Following the clustering process, we identified approximately 48,000 clusters. From this extensive collection, we randomly selected 1,000 representative clusters, ensuring that any remaining clusters containing these selected representatives were excluded. This resulted in a final training set comprising 45,000 unique clusters, each chosen to maximize the diversity and coverage of the structural space pertinent to our model’s training objectives.

### 4.2 Extract multimer inter-chain information using language model

PLMs for protein sequences have been widely recognized for their ability to effectively capture dependencies among residues, thereby providing meaningful feature embeddings [22, 24–26]. These models are typically trained using a masked language modeling (MLM) loss, where randomly masked (or mutated) amino-acid sequences are input into the network, which then attempts to recover the original sequences through self-attention. The attention weights in these models reflect the interactions between different residues [23] and can thus be naturally employed as initial pair features between residues.

Recent research [28] has confirmed that integrating chain relative positional encoding with pre-trained language models, and further pre-training the language model using group sequences derived from protein-protein interactions (PPIs) and complexes in the PDB, enhances the language model’s ability to extract inter-chain information. Previous work [29] has successfully applied this approach to sequence-based antibody structure prediction, achieving commendable performance.

We have further adapted this approach for TCR and pMHC systems, including using relevant data (e.g.,short peptides and sequence pairs in TCRStructDB we collected) for further pre-training and designing a masking strategy specifically for the CDRs of TCR. The ESM-2 model, which comprises 650 million parameters, was selected as the base model. Our PLM model, named ESM-PPI-TCR, employs the algorithm proposed in a previous study to distinguish different chains for further pre-training. We have also adapted the masking strategy of protein language models, tailoring it to paired TCR sequences and pMHC sequences. Detailed information and data for ESM-PPI-TCR are introduced in Supplementary C.2.

### 4.3 Sequence-based unliganded TCR & unbound pMHC prediction

We initiate our process by extracting the final sequence embeddings and all-layer pairwise attention weights from the pre-trained ESM-PPI model for each paired TCR chain and pMHC, while keeping the parameters fixed. For the model’s input, multiple chains are concatenated into a single sequence. To effectively distinguish between different chains, we utilize three distinct positional encoders: the multimer positional encoder, the relative positional encoder, and the chain relative positional encoder. These encoders are specifically designed to infuse additional positional information into the initial sequence embedding and pairwise representation, thereby enriching the model’s input data. Once the input sequence embedding is initialized, we employ the Evoformer-Single stack for iterative updates of single features and pair features. The Evoformer-Single module is identical to the one used in tFold-Ab [29], and is similar to the Pairformer module used in AlphaFold3 [19]. Following the update of sequence and pair representation for sequence pairs by the Evoformer-Single, we adopt an IPA-based structure module for atom-level 3D structure prediction.

While different TCRs exhibit variations in their CDRs, they share substantial structural similarities in other regions. Similarly, in pMHC structures, MHCs of the same type possess identical sequences and structures. Given the limited availability of TCR and pMHC structural data, training the model exclusively on TCR and pMHC could lead to overfitting. To counteract this, we initially pre-train the model using general monomers and multimers sourced from the PDB, followed by fine-tuning with TCR and pMHC structures. This approach effectively enhances the model’s performance. Considering the inconsistent data distribution of TCR and pMHC, we have trained two separate models. These models share identical architectures and are initialized using the same pre-trained model, differing only in the training data used.

### 4.4 TCR-pMHC complex prediction with representation-driven flexible docking module

While the approach of using a pre-trained language model combined with the Evoformer-Single structure module can proficiently predict the structures of unliganded TCR and unbound pMHC, it struggles when applied to TCR-pMHC complexes. This limitation primarily stems from the scarcity of paired TCR-pMHC sequence data, which is not suitable for further pre-training of the ESM-PPI-TCR. Despite the consistent binding pattern of pMHC, the conformations of the binding regions (TCR CDR3 and peptide) differ significantly, posing a challenge for the model to learn the intricacies of the binding region. Furthermore, the limited availability of TCR-pMHC structural data hampers the model’s ability to converge.

To better extract the interaction between TCR and pMHC, we have designed a feature fusion module. This module deviates from the traditional approach of using sequence features extracted from the language model as input. Instead, it utilizes features extracted from the pre-trained TCR and pMHC structure prediction models. For the initial sequence feature, we employ cross-attention [77] mechanisms to generate sequence embeddings for the TCR-pMHC complex. This mechanism allows each residue in one sequence to attend to all residues in the other sequence, thereby enhancing the interaction modeling. For the initial pairwise feature, we divide it into four segments for multimer structure prediction. The diagonal blocks are populated with pair features and initial coordinates derived from the TCR and pMHC components, while the off-diagonal segments are initialized with zero.

Following the feature fusion module, we input the initial single and pair features into a subsequent sub-network. This sub-network comprises 16 Evoformer-Single blocks and 8 structure modules, all sharing parameters. For the IPA-based structure module, we discovered that initializing with all-zero coordinates, as done in previous methods [17, 25, 29], is limited by the scarcity of training data and impedes the model’s convergence. Therefore, we initialized the structure module using the backbone structure of the unliganded TCR.

The final model outputs the coordinates of the TCR-pMHC complex. As the model considers the interaction between the TCR and pMHC during the modeling process, minor deformations occur at the interface of both components. Given that the pre-trained TCR and pMHC structure prediction models can predict unliganded TCRs and unbound pMHCs while extracting their respective representations, this module utilizes the outputs of these predictions as inputs. We refer to this prediction-based module as the representation-driven flexible docking module. This design choice is critical, as it enables the model to refine the binding site of the TCR-pMHC complex within a broader conformational context.

### 4.5 TCRStructDB: The largest sequence and structure databases of TCRs and pMHCs

To construct TCRStructDB, we collected and curated extensive sequence data for unliganded TCRs, unbound pMHCs, and TCR-pMHC complexes. The majority of TCR records were initially stored as raw sequencing files, which we processed to extract clean, annotated, and translated amino acid sequences in FASTA format. For pMHCs, we compiled data by pairing MHC alleles with peptides and associating pseudo-sequences of MHC with peptides to ensure direct and complete sequence pairings. For TCR-pMHC complexes, we gathered and organized data from known databases and relevant literature. Once we obtained the sequence data, we employed the tFold-TCR structural prediction method, recognized for its high precision and throughput, to simulate the structures of the TCR and pMHC libraries, as well as their complexed forms. This process involved the following steps:

- **Data Collection and Processing**: Raw sequencing files for TCRs and pMHCs were obtained from various public repositories and existing databases. These sequences were then annotated and translated into amino acid sequences, ensuring non-redundancy and accuracy.
- **Structural Prediction**: The tFold-TCR method was employed to predict the structures of the collected sequences. This included unliganded TCRs, unbound pMHCs, and TCR-pMHC complexes.
- **Database Construction**: The predicted structures and sequences were organized into TCRStructDB, a comprehensive database that includes both sequence and structural information.

This methodological approach enabled the creation of TCRStructDB, which not only provides a rich structural library but also facilitates the development of a vectorized database for representing TCR similarities. In the following sections, we will provide a detailed description of the collection and processing steps for each type of sequence data: TCR, pMHC, and TCR-pMHC complexes. This will include the specific sources of the data, the criteria for inclusion, and the methods used to ensure the accuracy and completeness of the sequences.

#### 4.5.1 TCR sequence data

We collected TCR sequence data from multiple sources, including existing single-cell TCR sequence databases, the 10x Genomics website [60], and raw sequencing data reported in various scientific literature. Notable TCR databases, such as the Observed T cell receptor Space (OTS) [78] and huardb [79], contain extensive repositories of sequence data enriched with metadata. During the curation process, we recorded relevant metadata alongside the structural sequences, including T cell subtypes, disease states, species, and other critical annotations. This comprehensive approach ensures that our dataset captures not only the genetic and structural diversity of TCRs but also contextualizes them within their biological and clinical environments, enabling a more nuanced analysis of TCR repertoire characteristics across different conditions and populations.

Earlier databases constructed from bulk sequencing, such as TCRdb [80] and immuneCODE [81], primarily contain TCR single-chain data. While our model (tFold-TCR) can simulate the structure of single chains, we believe that single-chain data alone cannot fully capture the complete structural dynamics of TCRs. Therefore, such data is not within the scope of our collection.

For files obtained from existing databases, we further process them to ensure they are formatted as full-length TCR amino acid sequences, which is essential for our analyses. For processing raw sequencing data, we employed a method similar to that used by the OTS database. We began by identifying relevant studies that utilized high-throughput sequencing technologies, such as 10x Genomics, to profile T cell receptor repertoires. SRA numbers from each study were used to retrieve FASTQ files via the SRA toolkit (version 2.9.2) [82]. The raw FASTQ files were downloaded and processed using the 10x Genomics CellRanger software (version 8.0.1) [83], which enabled the reconstruction of paired TCR sequences from single-cell data.

To ensure the integrity of the full-length TCR sequences, we utilized the ANARCI [84] tool to assign standardized numbering and extract the Fv region sequences necessary for structure prediction with the tFold-TCR. If any of the following issues were encountered: (1) missing fragments within the recorded full-length sequences; (2) ANARCI failing to identify the corresponding Fv region directly; (3) the identified Fv region lacking the CDR3 segment recorded in the original data; (4) the Fv region contains stop codons or ambiguous amino acids (represented by “*”), we employed the Stitchr [85] tool to reconstruct an appropriate full-length sequence. Stitchr uses available V and J gene data, CDR3 sequence information, and species metadata to generate a valid full-length TCR sequence. This reconstructed sequence undergoes further processing with ANARCI, which identifies a complete and reasonable Fv segment for subsequent analysis. Any sequences that cannot be processed by either Stitchr or ANARCI are removed from the dataset. Additionally, sequences generated using the Stitchr tool are annotated with a specific identifier in the database, allowing for easy differentiation from the original data.

#### 4.5.2 pMHC sequence data

To collect pMHC binding data, we utilized various widely recognized datasets, including those used for training and testing of the NetMHCpan-4.1 and NetMHCpan-4.0 methods [86], IEDB2016 [87], IC50test [88], Binarytest [89], MHC- I2020 [90], MHC-II2020 [90], and MHC-I2015 [91]. These datasets encompass binding affinity measurements and epitope information for different MHC-I and MHC-II molecules. For the Eluted Ligand (EL) single allele data, we ensure meticulous recording and annotation to maintain dataset integrity. Regarding the binding affinity data, affinity values are transformed into a standardized range (0 to 1) using the following formula:

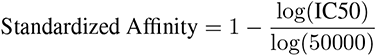

Thresholds for MHC classes were set based on binding affinities: For MHC-I, an IC50 value of less than 500 nM corresponds to a standardized affinity greater than 0.426; for MHC-II, an IC50 value of less than 1000 nM corresponds to a standardized affinity greater than 0.362 [90]. pMHC complexes with affinities greater than these thresholds are considered reasonable, indicating a likely biologically relevant binding interaction.

For multi-allele data, we utilize advanced models, specifically NetMHCpan [86], to perform precise annotation. Each peptide was assigned to its likely corresponding MHC allele by selecting the allele with the smallest EL rank score. Subsequently, we created a mapping table that links these allele names to their protein sequences using the IPD-MHC/HLA database [92].

To accommodate variations in allele naming conventions found in the original data, we manually curated relationships between different allele names and their aliases. For alleles not covered by the IPD database, such as certain mouse alleles, we consulted the UniProt [32] database to supplement their corresponding protein sequences. This process allowed us to construct complete pMHC sequences. Additionally, species annotations were standardized as “human”, “mouse”, or “other” to ensure consistent categorization across the dataset.

#### 4.5.3 TCR-pMHC sequence data

Our TCR-pMHC complex sequence data were collected from several sources:

- **3D Structure Databases**: This includes resources such as TCR3d [93] and STCRDab [37].
- **Affinity Databases**: Notably, we utilized the ALALS [63] database.
- **10x Antigen-Peptide Specific Stimulation Experiments** [60]: These experiments providing data on TCRs stimulated by specific antigens.
- **Existing TCR-Antigen Specificity Databases**: This category includes databases like VDJdb [49], McPAS-TCR [94], and the pan immune repertoire database (PIRD) [95].
- **Literature**: A limited set of data was extracted from relevant scientific publications, providing additional context and insights into TCR-pMHC interactions.

The collected data were processed using a standardized mapping and processing workflow for TCR and pMHC sequences, as previously described. This systematic approach allowed us to generate a comprehensive and consistent dataset of TCR-pMHC sequences, ensuring its suitability for downstream analysis.

## 5 Data Availability

Users can access our online platform, TCRStructDB, at https://ai4s.tencent.com/tcr. The original collected data for TCRStructDB is available upon reasonable request by contacting the corresponding author via email.

## 6 Code Availability

The source code, weights and inference scripts for the tFold models are available at https://github.com/TencentAI4S/tfold.

## Acknowledgments

We thank our colleagues at AI Lab and AI for Life Sciences Lab in Tencent for their encouragement and support.

## Author Contributions

Conceptualization: Y.Z., F.Wu., J.Y.; Methodology: F.Wu., Y.Z., Y.X.; Model development and software: F.Wu., Y.X., C.Q., and L.H.; Investigation and analysis: F.Wu., Y.X., Y.Z., and B.H.; Online platform development: Y.Z., Y.X., F.Wa., and Z.W.; Writing: F.Wu., Y.Z., Y.X., X.L., J.S., B.H., J.R., and J.Y.; Supervision: J.R. and J.Y.

## Appendix A Supplementary Figures

**Fig. A1:**
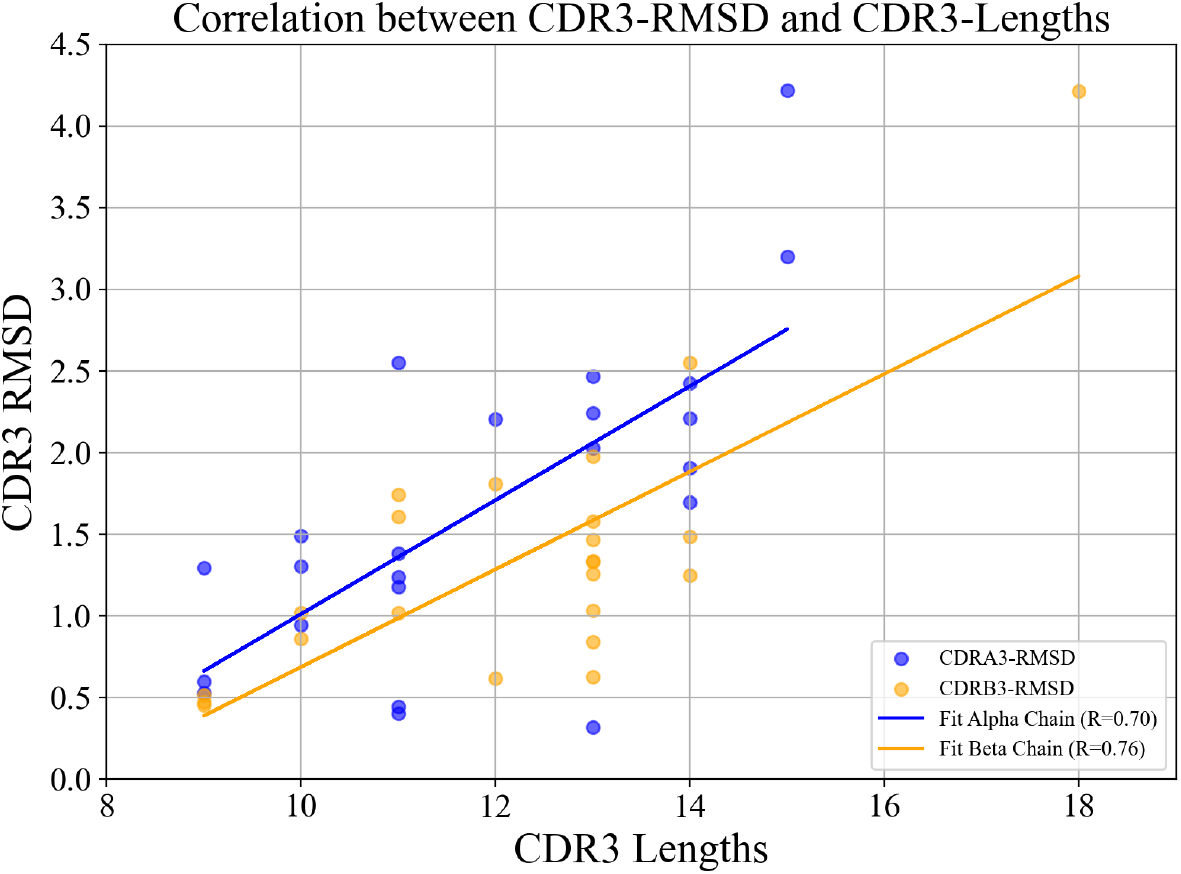
Analysis of RMSD values, derived from tFold-TCR predictions, against CDR lengths for CDR-A3 and CDR- B3, including linear fits and Pearson correlation coefficients to assess the strength of the relationships.

**Fig. A2:**
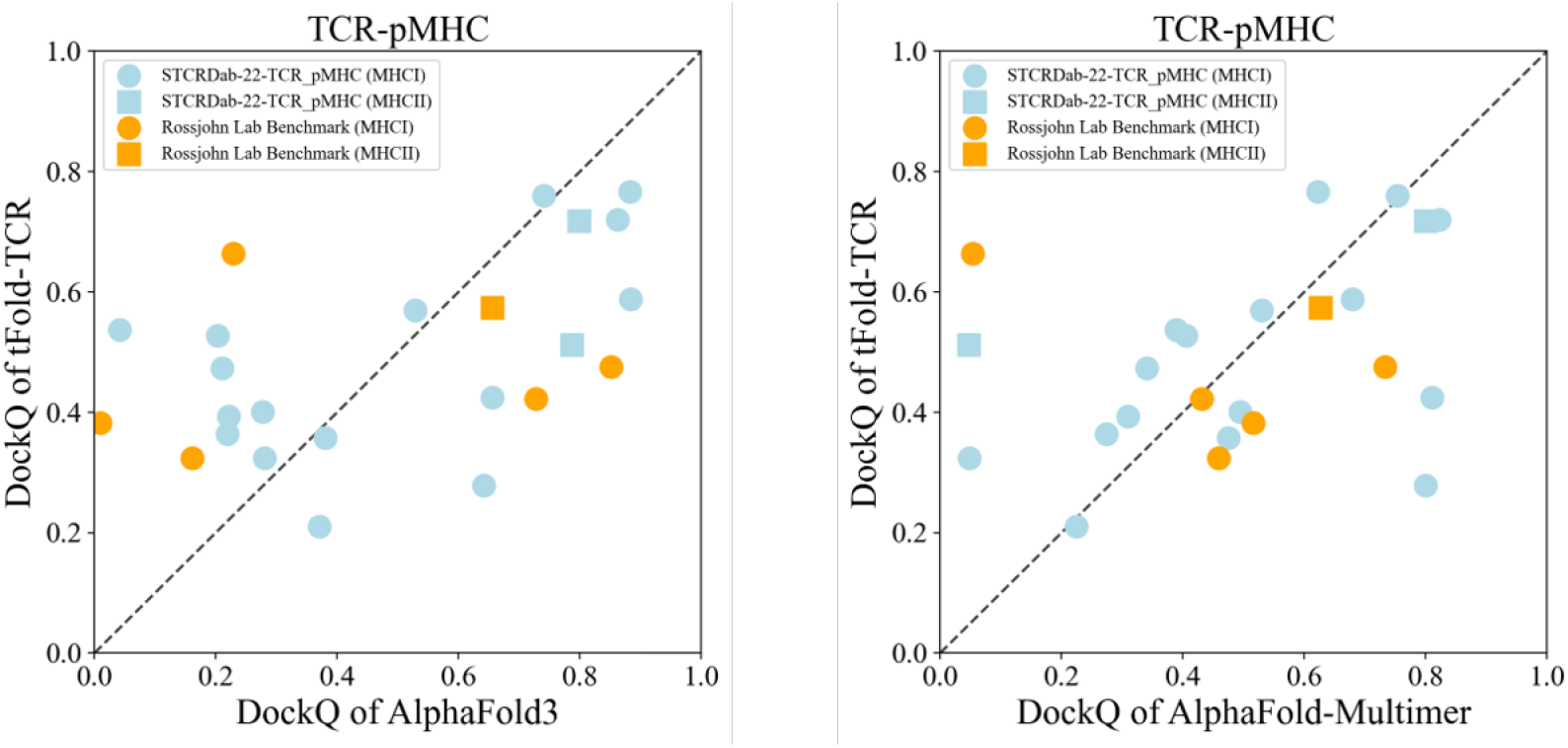
Comparison with tFold-TCR and other methods. Head-to-head comparison between tFold-TCR and AlphaFold- 3 (left), AlphaFold-Multimer (right) on STCRDab-22-TCR pMHC and Rossjohn Lab Benchmark.

**Fig. A3:**
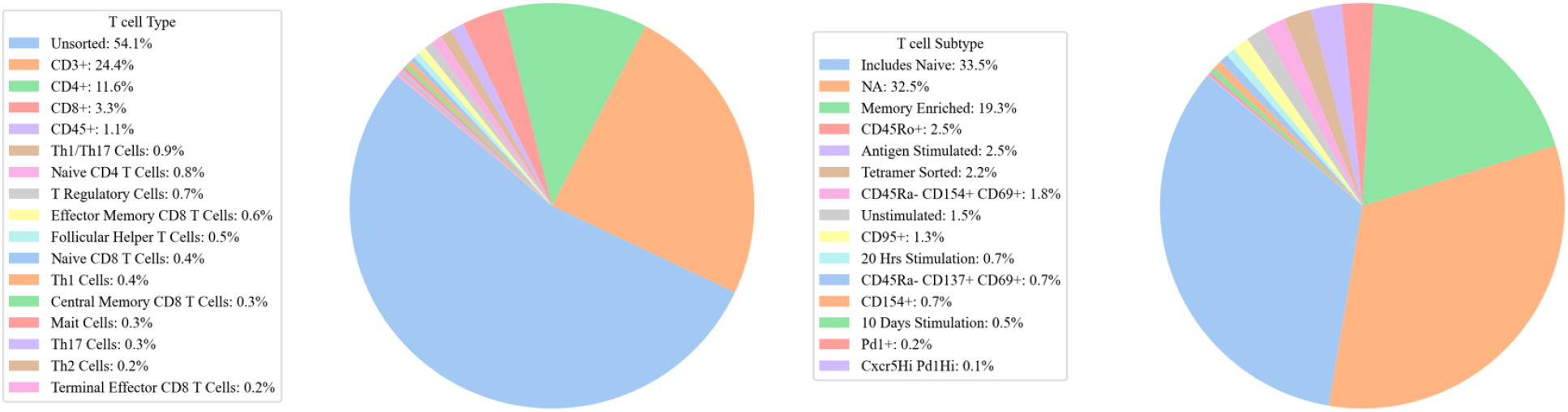
Metadata analysis of TCRStructDB. Pie charts illustrate the distribution of T cell type (left) and T cell subtype (right) within the database. Categories representing less than 0.1% are not shown.

**Fig. A4:**
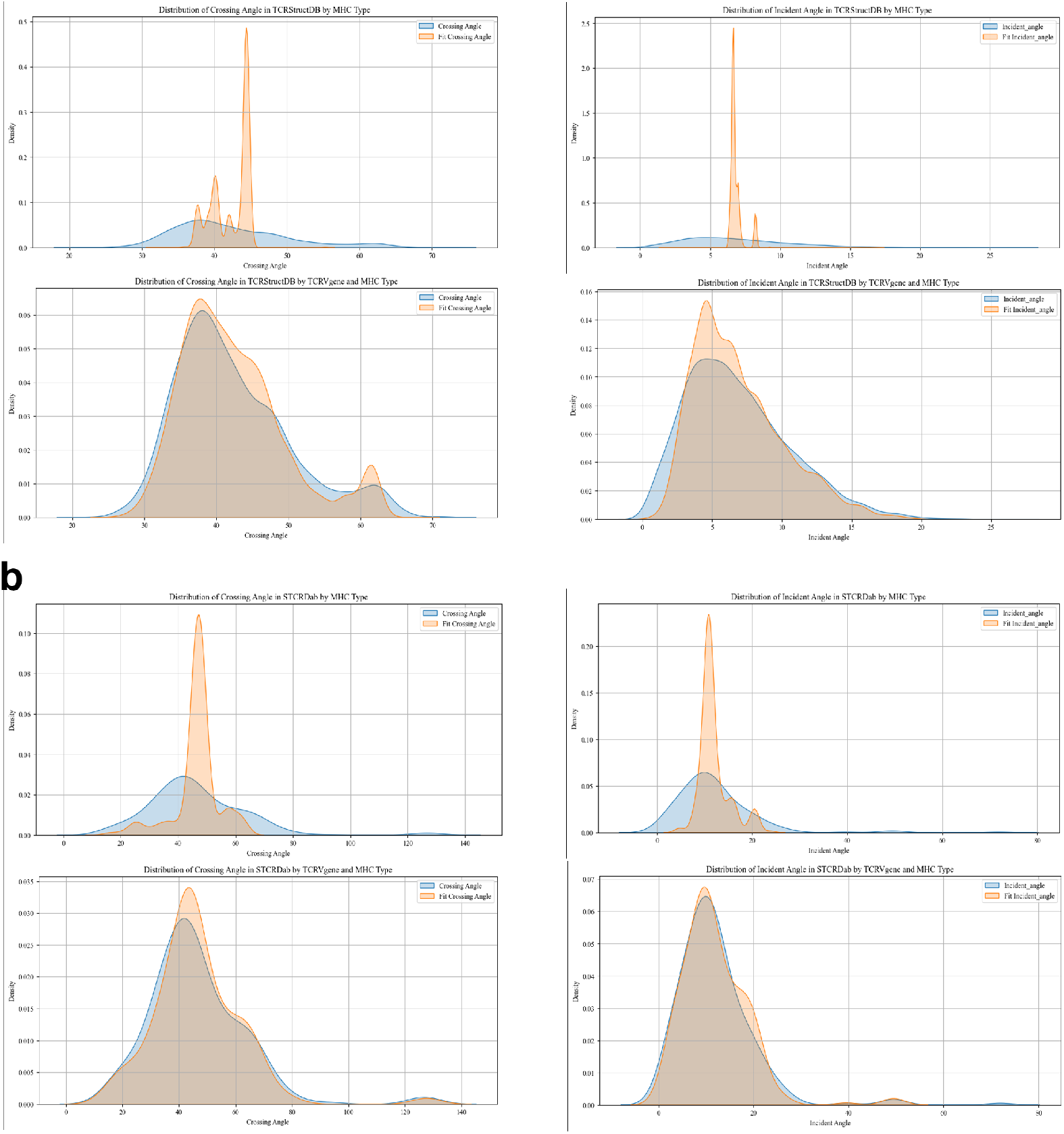
Single-Factor (MHC type) and multi-Factor (TCRV gene and MHC type) ANOVA Analysis of docking Angles in TCRStructDB and STCRDab. The distribution of docking angles (including crossing angle and incident angle) in the TCRStructDB (a) and STCRDab (b) databases. Single-factor and multi-factor ANOVA analyses were performed to demonstrate that docking angles are not solely determined by the MHC type. The single-factor model as an independent variable fails to adequately fit the distribution curves, whereas the multi-factor model provides a better fit.

## Appendix B Supplementary Tables

**Table B1:** Performance of unbound TCR structure prediction on the STCRDab-22-TCR benchmark. Backbone RMSD in different framework and CDR regions are reported. The TCR numbering scheme employed is IMGT. A-Fr indicates the Fr of A chain and A1-A3 indicate the CDRs of A chain. B-Fr indicates the Fr of B chain and B1-B3 indicate the CDRs of B chain.

| Method | A-Fr | A1 | A2 | A3 | B-Fr | B1 | B2 | B3 |
| --- | --- | --- | --- | --- | --- | --- | --- | --- |
| AlphaFold3 [19] | 0.635 | 0.860 | 0.735 | 1.802 | 0.497 | 0.545 | 0.401 | 1.509 |
| AlphaFold-Multimer [18] | 0.605 | 0.895 | 0.718 | 1.887 | 0.661 | 0.551 | 0.449 | 1.615 |
| ESMFold[25] | 0.717 | 1.183 | 0.910 | 2.266 | 0.606 | 0.576 | 0.572 | 1.781 |
| TCRModel2[21] | 0.569 | 0.789 | 0.633 | 1.774 | 0.481 | 0.502 | **0.384** | 1.520 |
| tFold-TCR | **0.566** | **0.736** | **0.560** | **1.665** | **0.469** | **0.442** | 0.441 | **1.352** |

**Table B2:** Performance and runtime of unbound TCR prediction on the STCRDab-22-TCR benchmark.

| Method | DockQ↑ | Fnat↑ | LRMS↓ | iRMS↓ | RMSD↓ | GDT↑ | Time↓ |
| --- | --- | --- | --- | --- | --- | --- | --- |
| AlphaFold3[19] | 0.773 | 0.760 | 2.047 | 1.224 | 0.877 | 0.928 | 60 s |
| AlphaFold-Multimer[18] | 0.785 | 0.776 | 1.816 | 1.186 | 0.873 | 0.931 | 6610 s |
| TCRModel2[21] | 0.794 | **0.781** | 1.795 | 1.120 | 0.763 | 0.948 | 262 s |
| tFold-TCR | **0.797** | 0.762 | **1.638** | **1.070** | **0.704** | **0.956** | 1.23 s |

**Table B3:** Performance of unbound pMHC prediction on the STCRDab-22-pMHC benchmark.

| Method | DockQ↑ | Fnat↑ | LRMS↓ | iRMS↓ | RMSD↓ | GDT↑ | Time↓ |
| --- | --- | --- | --- | --- | --- | --- | --- |
| AlphaFold3[19] | 0.926 | 0.882 | 1.111 | **0.441** | **0.655** | **0.960** | 70 s |
| AlphaFold-Multimer[18] | **0.927** | **0.886** | **0.998** | 0.462 | 0.714 | 0.953 | 9995 s |
| tFold-TCR | 0.908 | 0.839 | 1.193 | 0.476 | 0.684 | 0.959 | 2.04 s |

**Table B4:**
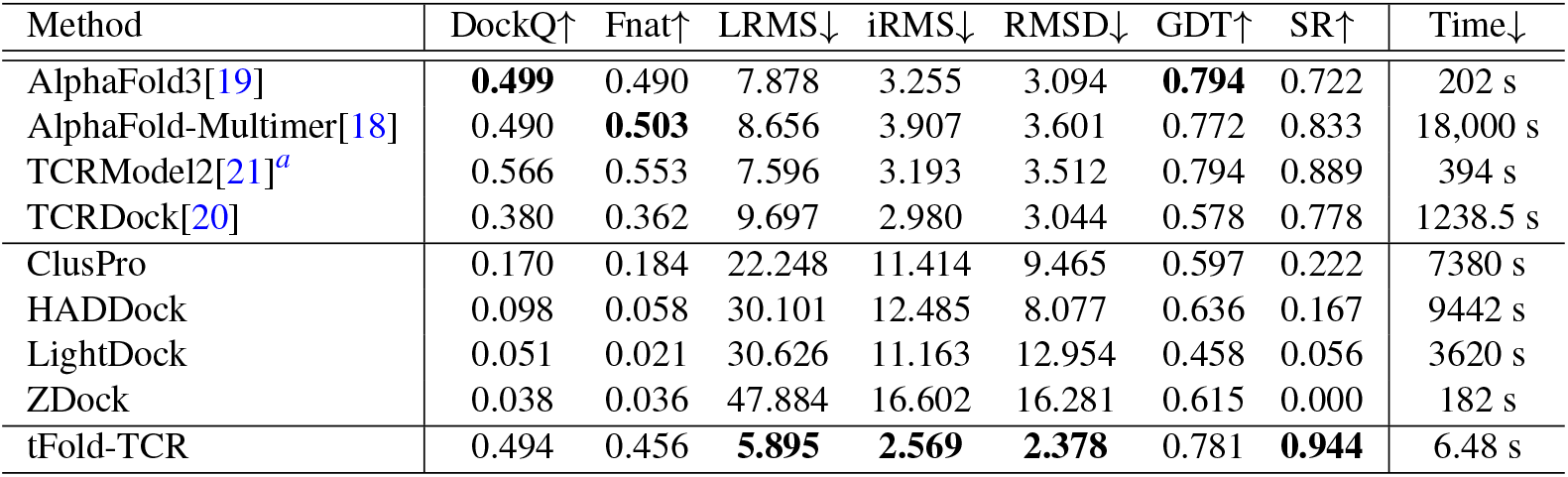
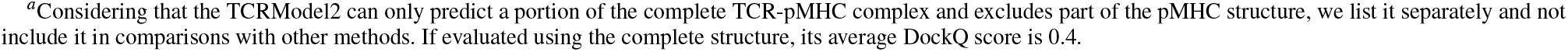
Performance of TCR-pMHC complex prediction on the STCRDab-22-TCR pMHC benchmark. SR denotes DockQ Success Rate as defined by DockQ algorithm.

| Method | DockQ↑ | Fnat↑ | LRMS↓ | iRMS↓ | RMSD↓ | GDT↑ | SR↑ | Time↓ |
| --- | --- | --- | --- | --- | --- | --- | --- | --- |
| AlphaFold3[19] | **0.499** | 0.490 | 7.878 | 3.255 | 3.094 | **0.794** | 0.722 | 202 s |
| AlphaFold-Multimer[18] | 0.490 | **0.503** | 8.656 | 3.907 | 3.601 | 0.772 | 0.833 | 18,000 s |
| TCRModel2[21]<sup>a</sup> | 0.566 | 0.553 | 7.596 | 3.193 | 3.512 | 0.794 | 0.889 | 394 s |
| TCRDock[20] | 0.380 | 0.362 | 9.697 | 2.980 | 3.044 | 0.578 | 0.778 | 1238.5 s |
| ClusPro | 0.170 | 0.184 | 22.248 | 11.414 | 9.465 | 0.597 | 0.222 | 7380 s |
| HADDOCK | 0.098 | 0.058 | 30.101 | 12.485 | 8.077 | 0.636 | 0.167 | 9442 s |
| LightDock | 0.051 | 0.021 | 30.626 | 11.163 | 12.954 | 0.458 | 0.056 | 3620 s |
| ZDock | 0.038 | 0.036 | 47.884 | 16.602 | 16.281 | 0.615 | 0.000 | 182 s |
| tFold-TCR | 0.494 | 0.456 | **5.895** | **2.569** | **2.378** | 0.781 | **0.944** | 6.48 s |

<sup>a</sup>Considering that the TCRModel2 can only predict a portion of the complete TCR-pMHC complex and excludes part of the pMHC structure, we list it separately and not include it in comparisons with other methods. If evaluated using the complete structure, its average DockQ score is 0.4.

**Table B5:** Performance of TCR-pMHC complex prediction on the Rossjohn Lab Benchmark. SR denotes DockQ Success Rate as defined by DockQ algorithm.

| Method | DockQ↑ | Fnat↑ | LRMS↓ | iRMS↓ | SR↑ |
| --- | --- | --- | --- | --- | --- |
| AlphaFold 3 | 0.439 | 0.407 | 15.52 | 4.819 | 0.500 |
| AlphaFold-Multimer | 0.470 | 0.468 | 8.844 | 4.669 | 0.833 |
| TCRDock | 0.385 | 0.327 | 9.634 | 3.852 | 0.667 |
| ClusPro | 0.011 | 0.000 | 60.684 | 19.23 | 0.000 |
| HADDOCK | 0.197 | 0.191 | 17.595 | 5.072 | 0.330 |
| LightDock | 0.009 | 0.000 | 59.212 | 17.956 | 0.000 |
| ZDock | 0.013 | 0.000 | 68.587 | 24.808 | 0.000 |
| tFold-TCR | **0.474** | **0.482** | **6.462** | **2.477** | **1.000** |

**Table B6:**
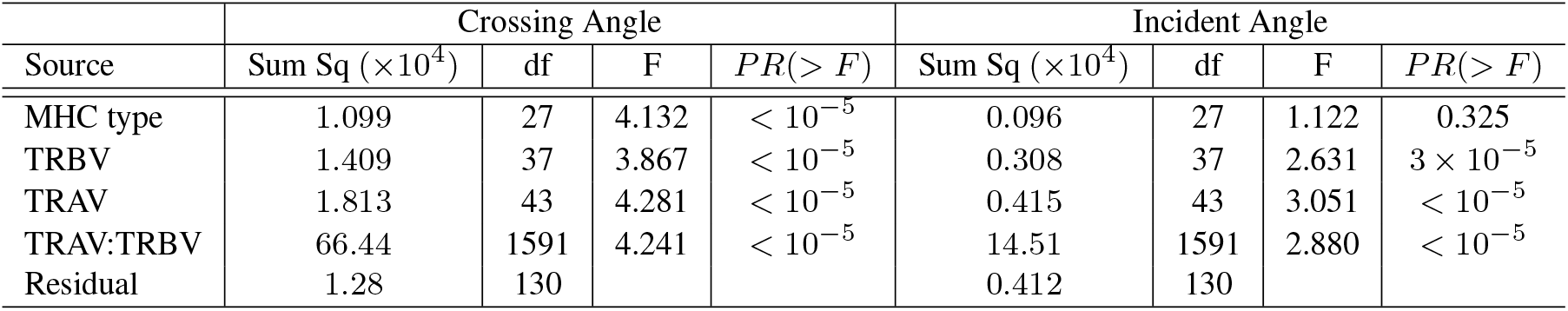
Summary of ANOVA for docking angles derived from STCRDab.

| Source | Crossing Angle |  |  |  | Incident Angle |  |  |  |
| --- | --- | --- | --- | --- | --- | --- | --- | --- |
| | Sum Sq ($\times 10^4$) | df | F | $PR(> F)$ | Sum Sq ($\times 10^4$) | df | F | $PR(> F)$ |
| MHC type | 1.099 | 27 | 4.132 | $< 10^{-5}$ | 0.096 | 27 | 1.122 | 0.325 |
| TRBV | 1.409 | 37 | 3.867 | $< 10^{-5}$ | 0.308 | 37 | 2.631 | $3 \times 10^{-5}$ |
| TRAV | 1.813 | 43 | 4.281 | $< 10^{-5}$ | 0.415 | 43 | 3.051 | $< 10^{-5}$ |
| TRAV:TRBV | 66.44 | 1591 | 4.241 | $< 10^{-5}$ | 14.51 | 1591 | 2.880 | $< 10^{-5}$ |
| Residual | 1.28 | 130 |  |  | 0.412 | 130 |  |  |

**Table B7:** Summary of ANOVA for docking angles derived from TCRStructDB.

| Source | Crossing Angle |  |  |  | Incident Angle |  |  |  |
| --- | --- | --- | --- | --- | --- | --- | --- | --- |
| | Sum Sq ($\times 10^4$) | df | F | $PR(> F)$ | Sum Sq ($\times 10^4$) | df | F | $PR(> F)$ |
| MHC type | 0.742 | 27 | 21.052 | $< 10^{-5}$ | 0.094 | 27 | 8.304 | $< 10^{-5}$ |
| TRBV | 0.423 | 53 | 6.112 | $< 10^{-5}$ | 0.236 | 53 | 10.579 | $< 10^{-5}$ |
| TRAV | 3.470 | 35 | 75.922 | $< 10^{-5}$ | 0.861 | 35 | 58.452 | $< 10^{-5}$ |
| TRAV:TRBV | 21.45 | 1855 | 8.856 | $< 10^{-5}$ | 6.400 | 1855 | 8.197 | $< 10^{-5}$ |
| Residual | 7.903 | 6053 |  |  | 2.547 | 6053 |  |  |

## Appendix C Supplementary Methods

### **C.1** Overview

In this section, we provide a comprehensive explanation of the methods and results described in the main text. First, we present the details of our ESM-PPI-TCR model in Section C.2. Next, in Section C.3, we describe how we modified the tFold-TCR model to make it suitable for structure-sequence co-design tasks and constructed a test set through computational means for evaluation. Following this, we provide a user guide for TCRStructDB online platform in Section C.4.

### **C.2** ESM-PPI-TCR

#### **C.2.1** Datasets

A combination of six datasets is used in the further pre-training for ESM-PPI:

- **UniRef50** [96] (March 2023 version): Approximately 60 million monomeric sequences. We randomly selected 4,000 sequences that have never appeared in the training and validation sets of ESM2 to ensure the validity of our validation set. These sequences were obtained by taking the difference set between UniRef50 and UniRef100 released in September 2021, and were used to construct the validation and test sets.
- **PDB** (January 2023 version): Comprising a total of 188,000 sequences of solved protein multimers in the Protein Data Bank. We selected 4,000 pairs of interacting chains from 4,000 complex structures released after 2022 to serve as the validation and test sets.
- **PPI**: A dataset compiling 1.3 million pairs of protein sequences known to potentially interact, gathered from various existing databases. We amalgamated pairs of interacting proteins from sources such as HINT [97], intACT [98], HIPPIE [99], prePPI [100], BioGRID [101], comPPI [102], and huMAP [103]. To address redundancy, we employed MMseqs2 [74] to filter out duplicated PPIs based on 100% sequence identity. Additionally, we filtered out interactions with low confidence scores in intACT (score lower than 0.3), comPPI (score lower than 0.3), and HIPPIE (quality lower than 0.63) to ensure higher data quality. After filtering, we selected 4,000 pairs for validation and test sets.
- **Antibody**: This dataset includes 1.5 million paired antibody sequences collected from the OAS [104]. Considering that the antibody’s CDR3 region can be affected by the antigen, we did not select any paired antibody as a validation or test set.
- **TCR-pMHC**: This dataset consists of all sequence pairs in TCRStructDB, constructed as part of this work.
- **Peptide**: A dataset gathered from PeptideAtlas [105], DBAASP [106], APD [107], SATpdb [108], and the aforementioned target-peptide pairs. The dataset was clustered at 50% sequence identity using MMseqs2 to guarantee sufficient structural diversity between training and validation sets, containing 0.6 million peptides. Sequences containing fewer than 10 amino acids were excluded to ensure the effectiveness of masked language modeling.

#### **C.2.2** Training details

The training process of ESM-PPI-TCR resembles that of traditional masked language models. During training, 15% of amino acids are randomly selected for masking: 80% of these are masked, 10% are substituted with a random residue, and 10% are left unchanged. To prevent data leakage in homomers, we ensure that the masking is uniform across all chains. For antibodies and TCRs, due to the predictability of amino acid types in the framework regions, we implement a strategy where residues in the CDRs have a 30% higher probability of being chosen for masking, while the framework regions remain unchanged. Given the short length of peptides, a 15% masking strategy may not provide sufficient information for effective learning; therefore, we increase the masking probability for peptides to 30%.

In the further pre-training phase, we executed a comprehensive 128,000 training steps, with each step comprising a batch of 128 samples. These samples varied, including individual sequences from UniRef50, pairs from multimeric PDB structures, PPI interaction pairs, paired antibody sequences, TCR-pMHC sequences, and peptides. The selection probability for these sample types was uniformly distributed.

We utilized the AdamW optimizer with hyperparameters *β*_1_ = 0.9, *β*_2_ = 0.999. The learning rate schedule started with a linear warm-up from 3e*−*6 to 3e*−*5 over the first 12,800 steps, after which it was maintained at 3e*−*5 for the rest of the training. The entire additional pre-training phase was executed on a cluster of 8 NVIDIA A100 GPUs, completing in roughly 30 hours. This configuration enabled us to scale our training process efficiently while optimizing the use of computational resources.

#### **C.2.3** Distinguishing TCR through ESM-PPI-TCR embeddings

The relationship between sequence similarity and functional or structural similarity is well-established in molecular biology. In this context, we aim for ESM-PPI-TCR to effectively differentiate between various TCR sequences.

To assess the model’s capability, we generated embeddings for TCRs in TCRStructDB. Each paired *αβ*TCR sequence was directly input into ESM-PPI-TCR, and the resulting embeddings for each chain were segmented according to their respective lengths. We visualized these embeddings using a t-SNE plot, where each data point corresponds to a paired *αβ*TCR sequence.

The V gene plays a pivotal role in shaping the Fv region of the TCR. As depicted in Fig. C5, the embeddings generated by ESM-PPI-TCR exhibit distinct distributions for sequences associated with different V genes, irrespective of their association with either the alpha or beta chain. This demonstrates that ESM-PPI-TCR effectively captures sequence-specific information, highlighting its capability to differentiate between various TCR types. To further explore whether ESM-PPI-TCR can distinguish between different antigenic features, we collected TCRs from TCRStructDB that bind to diverse epitopes. We selected the ten antigens with the highest number of associated TCRs and generated embeddings using ESM-PPI-TCR, as shown in Fig. C6. However, since ESM-PPI-TCR has not been fine-tuned for downstream binding/non-binding prediction tasks, it cannot accurately predict the interactions between TCRs and peptides. This also underscores the inherent complexity of this challenge.

**Fig. C5:**
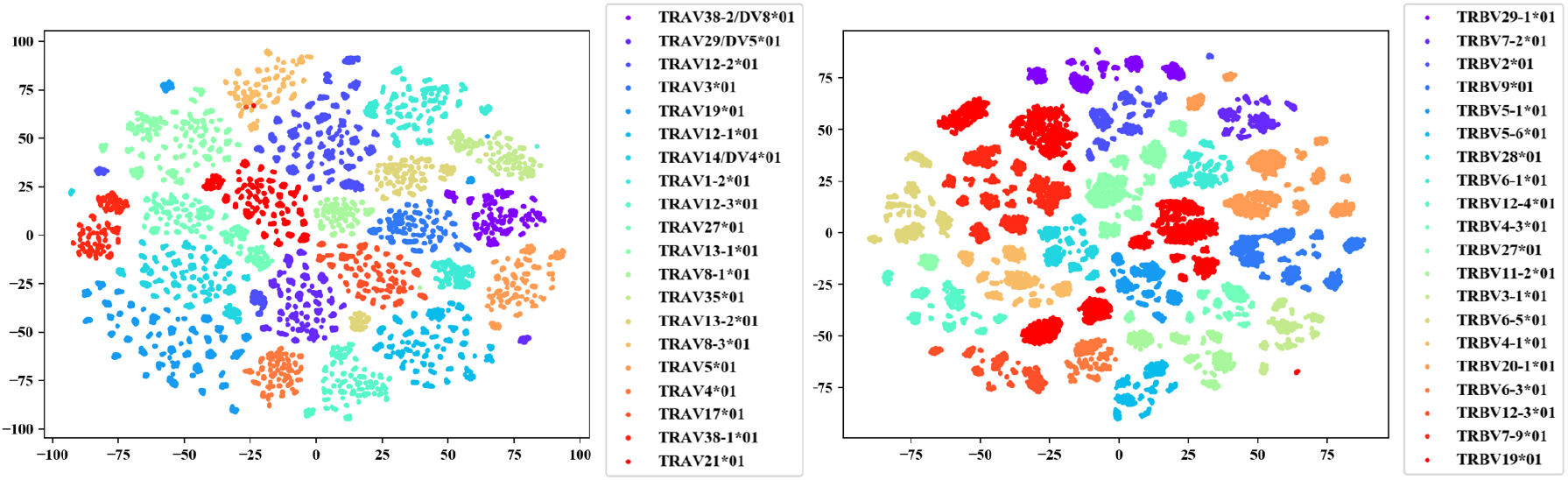
t-SNE visualization of ESM-PPI-TCR sequence embeddings for TCR alpha chain (left) and beta chain (right) colored by V gene categories.

**Fig. C6:**
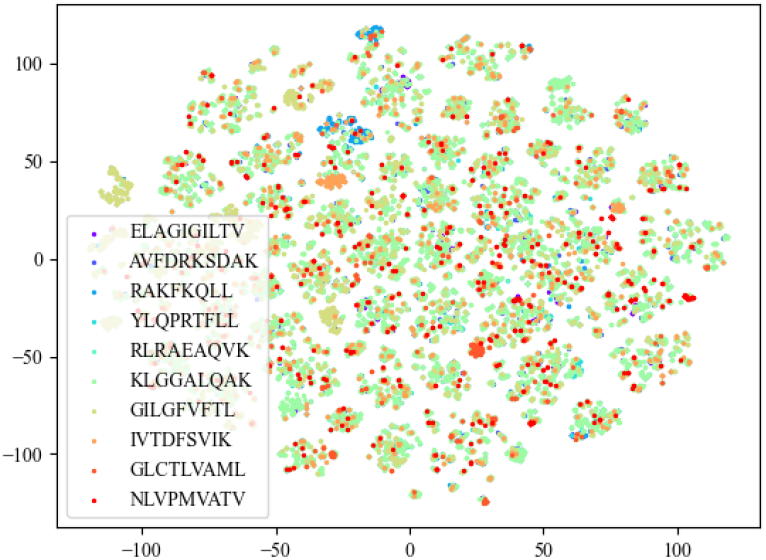
t-SNE visualization of ESM-PPI-TCR sequence embeddings colored by antigen categories.

### **C.3** CDRs design with tFold-TCR

#### **C.3.1** tFold-TCR for structure-sequence co-design

The structural prediction task [17–19] involves predicting the 3D structure of a protein based on its amino acid sequence. In contrast, sequence recovery [109–111] focuses on reconstructing the amino acid sequence from a given protein backbone structure. Additionally, structure-sequence co-design [29, 68] entails predicting the overall structure of a protein while recovering masked portions of the sequence, thereby integrating both structural and sequence information in a unified framework.

We implement structure-sequence co-design through a multi-task learning approach. Specifically, during the training process, we replace the amino acid types in the CDRs of TCRs with a *<*Mask*>* token with a probability of 30%. Importantly, we do not mask other regions of the TCR or the pMHC sequences. These masked TCR sequences, along with the corresponding antigen sequences, are then utilized as input for the model. To enhance our representation-driven flexible docking module, we introduce an additional auxiliary head, implemented as feed-forward layers, to predict the masked amino acid types. Ultimately, the original unmasked TCR sequences provide supervision for the TCR design head, while the experimentally determined structures serve as supervision for the complex conformation prediction head.

Through this multi-task learning training mechanism, we aim for tFold-TCR to consider the interactions between TCRs and pMHC during the prediction process. This approach specifically emphasizes the interactions between TCRs and peptides, thereby encouraging the model to learn to generate new TCR sequences that are not only reasonable but also structurally similar to reference TCR structures.

#### **C.3.2** Benchmark and evaluation criteria

To evaluate the ability of tFold-TCR in designing TCRs, we constructed a test set named STCRDab-22-Design. This test set comprises 18 sequence pairs, consistent with STCRDab-22-TCR pMHC, but with the corresponding CDRs in the input TCR sequences replaced by *<*Mask*>* tokens.

Given the complexity of wet-lab validation for TCR design tasks, we employ an in silico evaluation metric: Amino Acid Recovery (AAR). AAR is defined as the overlap ratio between the generated sequence and the known binding TCR, ranging from 0 to 1. Higher AAR values indicate a greater similarity between the recovered TCR and the real TCR, thereby reflecting a stronger TCR design capability.

#### **C.3.3** In silico evaluation of CDRs design

Since CDR-A3 and CDR-B3 are the primary regions where *αβ*TCRs bind to pMHC, and are crucial for determining the specificity and affinity of the TCR, we focus our evaluation on the design capabilities for these two regions.

Table. C8 presents the CDRs design performance of tFold-TCR in the STCRDab-22 benchmark. The average AAR for CDR A3 reached 0.332, while CDR B3 achieved an AAR of 0.407. Although the sequence design capabilities of tFold-TCR have not yet been validated through wet lab experiments, in silico evaluations demonstrate the potential of using tFold-TCR to design TCR antibodies and optimize TCR-peptide affinity.

Fig. C7 illustrates the workflow and performance of tFold-TCR in designing the TCR for 7t2b. Although the designed CDR-B3 region does not fully align with the actual structure, it achieved a CDR-B3 recovery rate of 0.7. This indicates that tFold-TCR is capable of generating TCR designs that are reasonably close to the target structure, demonstrating its effectiveness in TCR design tasks.

**Table C8:** CDRs deisgn accuracy on the STCRDab-22 benchmark. Evaluated by AAR.

| Method | A3 | B3 |
| --- | --- | --- |
| tFold-TCR | 0.332 | 0.407 |

**Fig. C7:**
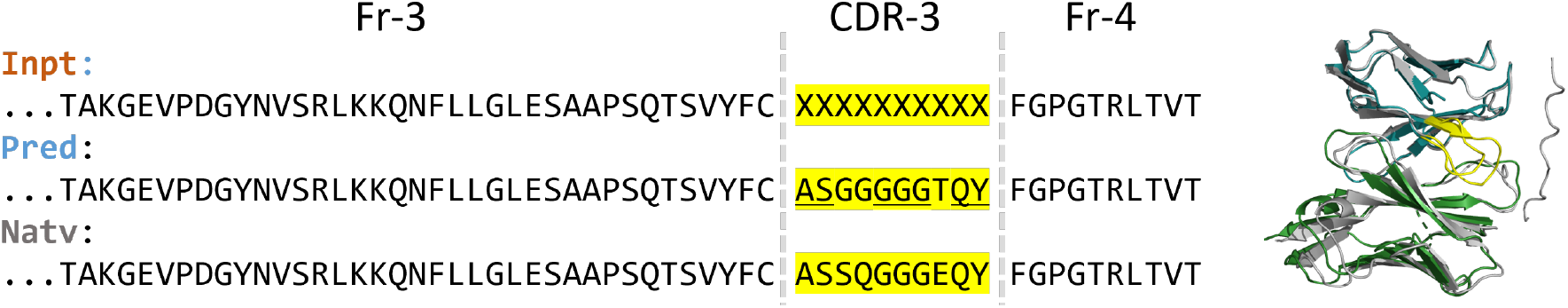
Comparison of our predicted structures and CDR-B3 amino acid type for an TCR target for 7t2b (blue for beta chain, green for alpha chain) with their respective experimental structures (gray). The CDR-B3 region highlight by yellow. tFold-TCR achieve 0.7 AAR for this case.

### **C.4** Utilization guide for TCRStructDB online platform

TCRStructDB oneline platform offers a suite of search and analytical web applications, which are categorized into three main functional groups: (1) Structure prediction, which corresponds to the “Structure prediction” interface; (2) Binding retrieval, associated with the “Interaction Query” interface; and (3) Similarity retrieval, linked to the “Similarity Search” interface.

The “Structure Prediction” web portal features tFold-TCR for predicting TCR-pMHC complexes, paired TCRs, pMHCs, and individual TCR or MHC structures. Based on the input chains, the portal provides the corresponding prediction functionality. For instance, if all five chains are inputted, the web server conducts a comprehensive TCR-pMHC structure prediction. Conversely, if only the paired chains of a TCR are provided, an unliganded TCR structure prediction is performed (refer to Table. C9 for more details).

A similar design is employed for the “Query Binding” and “Search Similar” web pages. The binding retrieval function aims to provide known (experimentally verified) binding items corresponding to the input. For instance, if paired TCR chains are provided, TCRStructDB will return the pMHCs known to bind to the input TCR, along with the corresponding TCR-pMHC complexes. For the similarity retrieval function, TCRStructDB enables users to find items similar to their inputs based on either sequence or structural characteristics. For example, if the input consists of paired chains of unliganded TCR, the server will return a set of TCRs with sequences or structures analogous to those of the input (details are shown in Table. C10 and Table. C11).

Furthermore, when users apply the search application, in addition to the provided sequence and structure information, TCRStructDB also offers corresponding metadata. This metadata includes detailed information related to TCR data such as Accession, Author, Disease, Cancer Type, Age, Species, Strain, TCell Source, Tcell Subtype, Treatment, ANARCI number, and more.

**Table C9:** Utilization Guide for the Structure Prediction interactive webpage.

| Input | Webpage | Call-to-action button | Output |
| --- | --- | --- | --- |
| TCR $\alpha\beta$ or $\gamma\delta$ chains | Structure Prediction | RUN | The complex structure of unliganded TCR |
| peptide and MHC chains | Structure Prediction | RUN | The complex structure of peptide-MHC |
| MHC chains | Structure Prediction | RUN | The complex structure of MHC |

**Table C10:** Utilization Guide for the Interaction Query interactive webpage.

| Input | Webpage | Call-to-action button | Output |
| --- | --- | --- | --- |
| TCR $\alpha\beta$ or $\gamma\delta$ chains | Interaction Query | Search | The pMHCs can bind to the input TCRs, structures of these TCR-pMHCs, along with relevant associated information. |
| peptide and MHC chains | Interaction Query | Search | TCRs can bind to the input pMHC, structures of these TCR-pMHCs, along with relevant associated information. |
| MHC chains | Interaction Query | Search | Peptides can be presented by the input MHC, structures of these pMHC, and relevant associated information. |
| peptide | Interaction Query | Search | MHCs can present the input peptide, structures of these pMHC, and relevant associated information. |

**Table C11:** Utilization Guide for the Similarity Search interactive webpage.

| Input | Webpage | Call-to-action button | Output |
| --- | --- | --- | --- |
| TCR $\alpha\beta$ or $\gamma\delta$ chains | Similarity Search | Search | TCRs with similar structures and sequences to the given input TCR |
| peptide and MHC chains | Similarity Search | Search | Peptides similar to the input peptide, presented by the same MHC, along with relevant associated information. |
| MHC chains | Similarity Search | Search | Similar peptides to the input peptide |

